# KPop: Accurate and scalable comparative analysis of microbial genomes by sequence embeddings

**DOI:** 10.1101/2022.06.22.497172

**Authors:** Xavier Didelot, Paolo Ribeca

## Abstract

The recent explosion in the amount of available sequencing data challenges existing analysis techniques. Here we introduce KPop, a novel versatile method based on full *k*-mer spectra and dataset-specific transformations, through which thousands of assembled or unassembled microbial genomes can be quickly compared. Unlike minimizer-based methods that produce distances and have lower resolution, KPop is able to accurately map sequences onto a low-dimensional space. Extensive validation on simulated and real-life viral and bacterial datasets shows that KPop can correctly separate sequences at both species and sub-species levels even when the overall genomic diversity is low. KPop also rapidly identifies related sequences and systematically outperforms minimizer-based methods. KPop’s code is open-source and available on GitHub at https://github.com/PaoloRibeca/KPop.

## Background

The last 20 years have seen such a revolution in high-throughput sequencing technology that it is now possible to sequence whole microbial genomes quickly and at low cost, enabling their use for a wide range of applications in the epidemiology of infectious diseases [1, 2, 3, 4]. Consequently, many studies have recently been carried out comparing hundreds or thousands of microbial genomes within a population of interest, which can be put in context with the very large numbers of genomes available in publicly accessible databases such as GenBank [5], BIGSdb [6] and EnteroBase [7]. However, the bioinformatic process needed to fully compare microbial genomes remains complex and laborious.

The first bioinformatic step typically involves the analysis of the raw sequencing reads. It can be performed by first aligning the reads to a reference genome, for example using `bwa` [8], Bowtie [9], or GEM [10], followed by calling variants relative to the reference, for example using FreeBayes [11]. Alternatively, analysis can be done without the use of a reference genome, for example with Velvet [12] or SPAdes [13], in which case the process is known as *de noυo* assembly [14]. Neither of these two approaches is fully satisfactory. A reference-based approach requires a predetermined reference genome, the choice of which becomes of paramount importance. Only genomic regions within this reference can be compared, which creates a bias in the analysis and prevents pan-genome analysis [15]. Assembling the genomes *de noυo* does not suffer from these difficulties, but requires an additional step to align the assembled genomes against each other. A full alignment is typically expensive beyond a handful of genomes and sometimes challenging if the sequences have evolved sufficiently, although some tools such as progressiveMauve [16] or Mugsy [17] might prove adequate in this scenario. Therefore, alignment is often performed only for core regions or genes, for example using a cgMLST scheme [18], or by aligning each genome against a reference, for example with MUMmer [19], which reintroduces the need for a reference and the loss of information on the accessory genome.

Here we propose a novel methodology for the comparison of large numbers of microbial genomes, that does not require assembly or alignment. Our approach is based on *k*-mers, which are nucleotide sequences of short length *k* found in the genomes. The use of *k*-mers in microbial genomics has been important before, especially for studies interested in quantifying the variability of the pan-genome [20, 21], including genome-wide association mapping studies that need to capture evolution in both core and accessory genome [22, 23].

Our strategy is conceptually different from most other comparable methods based on *k*-mers, which can be broadly divided into the following categories:

(1) Methods using specific, possibly very long *k*-mers to detect the presence or absence of specific sequences (as in Mykrobe [24]).
(2) “Binning” methods using all *k*-mers with a low value of *k* = 4 or 5 and relying on either machine learning to classify sequences (as in CONCOCT [25]) or on a dimensional reduction to cluster and visualize sequences in an abstract small-dimensional space (such as [26, 27]).
(3) Methods using simplified signatures (“sketches”) based on some dataset-independent choices (for instance, *k*-mer “minimizers” as in `mash` [28], `sourmash` [29] or PopPUNK [30]) to estimate genomic distances and summarise the relationships between sequences.

Instead, we use a full *k*-mer spectrum with a value of *k* typically ranging from 10 to 12 for bacterial species. This means that all the *k*-mers present in a genome and their frequencies contribute to the analysis, as opposed to MinHash-based methods that consider a small random sample of minimizing *k*-mers to improve efficiency [28] and often discard frequency information. For classification purposes, this is more robust than methods in category (1) above since it does not rely on long *k*-mers which can be disrupted by mutations, but it is still accurate because we consider a full spectrum. With respect to methods in category (2), using longer *k*-mers requires a more sophisticated algorithmic implementation due to the much larger size of the resulting *k*-mer spectra, but it also allows us to achieve a better precision based on more specific sequence matches. As for methods in category (3), we too use a transformation on *k*-mer spectra to reduce their dimensionality, but one that is tailored to the content of the dataset under analysis instead of arbitrarily relying on the use of minimizers [31]. Our method also produces a representation of each sample as a vector embedded in a reduced-dimension space (see Methods). This can be stored for later downstream analysis and used, for instance, to compute distances or implement classification workflows based on ML or AI algorithms. It should also be noted that methods in category (3), while arguably being the most popular at the moment due to the scalability they provide even when used directly on unassembled raw reads as in the case of `sourmash` [29], were never designed to go beyond categorisation at the species level; for instance, they may struggle with the comparison of closely related genomes [28], which explains attempts to make them more accurate [30].

In contrast, the methodology we describe here allows the comparison of large numbers of genomes with high resolution, be they assembled or not, based on the full set of *k*-mers present in all the sequencing reads or in some specific subset of them. To demonstrate the accuracy of our approach, we apply it to several simulated datasets for which the true relationships between genomes are known, and show that it can correctly classify sequences into lineages and rapidly identify related genomes. We also demonstrate the usefulness of our method on several real-life datasets, in the case of both viral and bacterial pathogens.

## Results

### General overview

Let us consider that we want to analyse a *dataset*, which is a collection of *samples*. In turn, each sample will usually consist of one or more files generated with highthroughput sequencing techniques, and associated metadata. Alternatively, the sample might be one or more genomic sequences previously assembled from the sequencing data. Input sequences can therefore be either NGS FASTQ read files or FASTA files containing assembled sequences.

The overall workflow we use and describe in this paper is depicted in Figure 1 and can be summarised in three steps:

**Figure 1.**
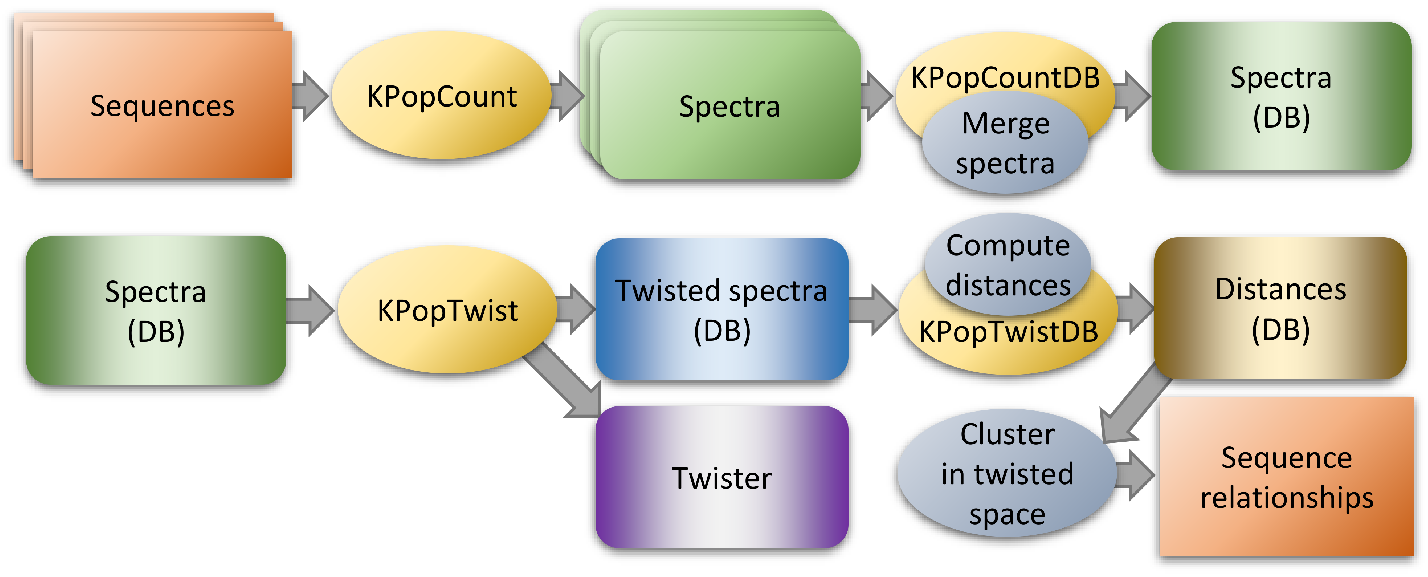
Basic workflow. Full *k*-mer spectra are deduced from input sequences and, after processing through KPopCount and `KPopCountDB`, stored into a binary database. These spectra can then be twisted to a reduced-dimensional representation thanks to `KPopTwist`; the resulting twisted vectors can be used to establish relations between sequences through their distances using `KPopTwistDB` or directly with other classification methods (not illustrated).

1. First, we consider the unbiased frequencies of *all k*-mers for a given and sufficiently large value of *k*, after suitable denoising and rescaling have been optionally performed.
2. Second, we determine a transformation (based on Correspondence Analysis, see Methods section) that performs a dimensional reduction step and is *optimised for the dataset at hand*. We call this transformation the “twister”; and we call “twisted” any *k*-mer spectrum to which the transformation has been applied. Computing the twister involves full *k*-mer spectra and is relatively compute- and resource-intensive. However, the optimisation of the twister could be performed on a representative subset of the sequences or after preclustering, thus reducing the final number of dimensions of the twisted spectra and the corresponding computational complexity. In practice, we found that this simplification was not needed for any of the examples shown below, as for most applications in microbial genomics the dimension of the twister is constrained by the number of taxonomic lineages, which is usually small. The twister can then be stored for future use (for instance, to perform clustering and/or classification). Once the optimal twister has been determined, its application to novel samples to twist their *k*-mer spectra takes a very little amount of computing time, which is determined by the dimension of the twister and independent of the sample.
3. Third, from the twisted sequences, classifiers can be built and relations between samples established. This can happen either via the computation of distances in spectral or twisted space or directly, for instance with methods such as decision trees [32] or random forests [33].

In general, the `KPop` method presented here has several desirable features. It produces unbiased (database-independent) spectra, that can be re-used in the future no matter which sequences are known at a given point in time. If novel sequences that are very different from what was already present in the database become known, one just has to regenerate the twister at step (2) and re-compute reduced-dimensional signatures from the spectra, which is quick and straightforward. When comparing strains of micro-organisms, our method can automatically use information about the full genome rather than just a set of core genes, which represents a substantial advantage compared to alignment-based methods. For instance, `KPop` is able to automatically take into account the presence of novel plasmids carrying antimicrobial resistance genes or other virulence factors even when the exact sequence for such genetic elements is not known or when the bacterial strains do not differ in their chromosomal sequences. On the other hand, by performing a suitable pre-processing of the data (see Methods), it is possible to focus on specific regions of the genome whenever needed. Finally, as demonstrated by one of the examples of application below, classifiers based on `KPop` are largely insensitive to recombination and other mechanisms that can obscure clonal relationships [34].

Many different variations on this general strategy can be implemented, especially if one provides the different computational blocks listed above as modular units that can be freely combined into more complex workflows. A detailed description of the protocols used in the rest of the paper, including commands to reproduce the examples, is available in the Methods section or online [35]. Unless otherwise stated, the tests mentioned in the text were performed on an Intel Xeon CPU E5-2680 v4 clocked at 2.40GHz, of which we only used 16/56 logical hyperthreads and up to 256GB of RAM. While for this paper we were not interested in an exact measurement of the time spent in each example, we do provide approximate timings.

### Classification at the bacterial species level

We followed the procedure illustrated in Figure 2 to perform classification using `KPop`. First we wanted to test the ability of this classifier to discriminate between bacterial species. We selected the 20 most highly represented bacterial species from a recent curated and searchable snapshot of bacterial whole genome sequences [36], namely *Salmonella enterica, Escherichia coli, Streptococcus pneumoniae, Mycobacterium tuberculosis, Staphylococcus aureus, Campylobacter jejuni, Listeria monocytogenes, Neisseria meningitidis, Streptococcus pyogenes, Clostridioides difficile, Klebsiella pneumoniae, Streptococcus agalactiae, Campylobacter coli, Neisseria gonorrhoeae, Enterococcus faecium, Pseudomonas aeruginosa, Vibrio cholerae, Acinetobacter baumannii, Mycobacteroides abscessus* and *Legionella pneumophila*. From each of these 20 species, we selected 40 genomes at random. The short read sequences of half of these genomes were used to train the classifier and the other half was used to test it. With `KPop`, there was only one genome out of 400 test genomes for which the classification was not the one previously reported (SRR5025480, previously reported as *Salmonella enterica* but classified as *Vibrio cholerae*). When the classification was performed with `sourmash tax` at default parameters, this genome was unclassified; there was also one misclassified genome (ERR1541368) and two other unclassified genomes (SRR5990258 and SRR8101476). Changing the value of *k* for `sourmash tax` from 31 to 21 resulted in one more genome (SRR5990258) becoming correctly classified, but ERR1541368 remained misclassified and SRR8101476 remained unclassified.

**Figure 2.**
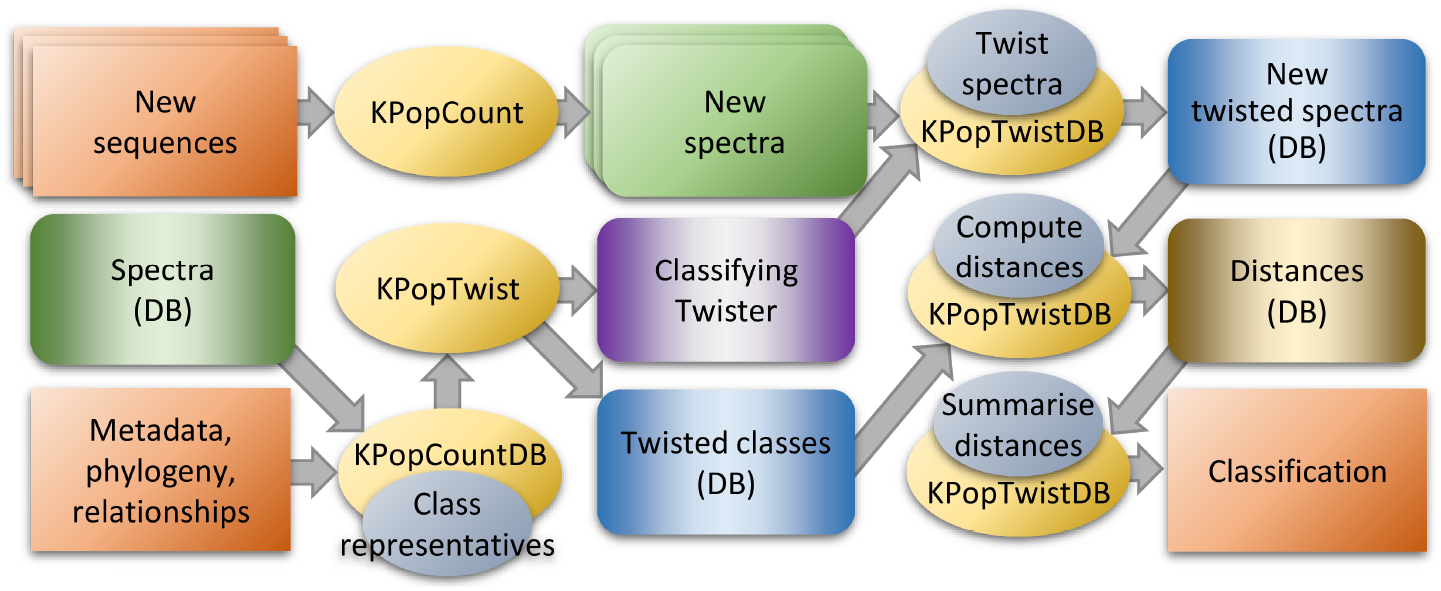
Classification workflow. **Lower left corner:** One starts with a set of spectra that have been computed according to the upper part of the basic workflow (Figure 1). We also assume that a suitable categorisation of such spectra into classes can be inferred from available metadata. One then computes representative spectra for each class, by generating with `KPopCountDB` a linear combination of the normalised spectra for all the sequences belonging to the class. The representative spectra are then twisted with `KPopTwist`. That returns a classifying twister (mauve box) and the twisted spectra for the representatives of each class (blue box). **Upper row:** Full *k*-mer spectra are computed with KPopCount for new sequences that we would like to classify; through `KPopTwistDB` they are twisted according to the transformation determined at the previous step, and stored into a database of twisted spectra. **Lower right**: Distances between the twisted spectra for the new sequences and the twisted spectra for class representatives are computed and summarised with `KPopTwistDB`. The name of the closest class representative is considered to be the correct classification. More class representatives, up to a set number of closest neighbours, can be considered to provide estimates of the likelihood of the classification and/or to implement *k*-NN classification schemes. **Not illustrated:** Alternatively, classifiers that do not imply computation of distances (for instance, using decision trees [32] or random forests [33]) can also be implemented if one starts directly from the twisted spectra for the *training* sequences and their classification as deduced from associated metadata.

Classification at the bacterial species level is relatively easy due to the high number of differences in both the core and accessory genomes when comparing genomes from different species. We note that the `KPop` classification was perfect even for closely related species that are known to recombine between each other, for example *Campylobacter jejuni* and *C. coli* [37]. The next examples test the ability of `KPop` to perform classification at a finer resolution.

### Classification of simulated tuberculosis genomes

We simulated a dataset of 1000 genomes of *Mycobacterium tuberculosis*. First a genealogy was generated with samples taken between year 2000 and 2022, at random from one of ten lineages that emerged in year 1900 and last shared a common ancestor around year 1500 (cf Figure 3A). The ten lineages were sampled with the same probability, so that the number of samples from each lineage was relatively constant, ranging between 75 and 130. Genomes of length 4.4 Mbp were then generated by mutation on the branches of this tree, with a substitution rate of 1.1 *×* 10^*−*7^ per year per site [38] being applied to the reference genome H37Rv [39]. Within each lineage, a randomly selected half of the genomes was used to train the classifier and the remaining half was used to test its accuracy.

**Figure 3.**
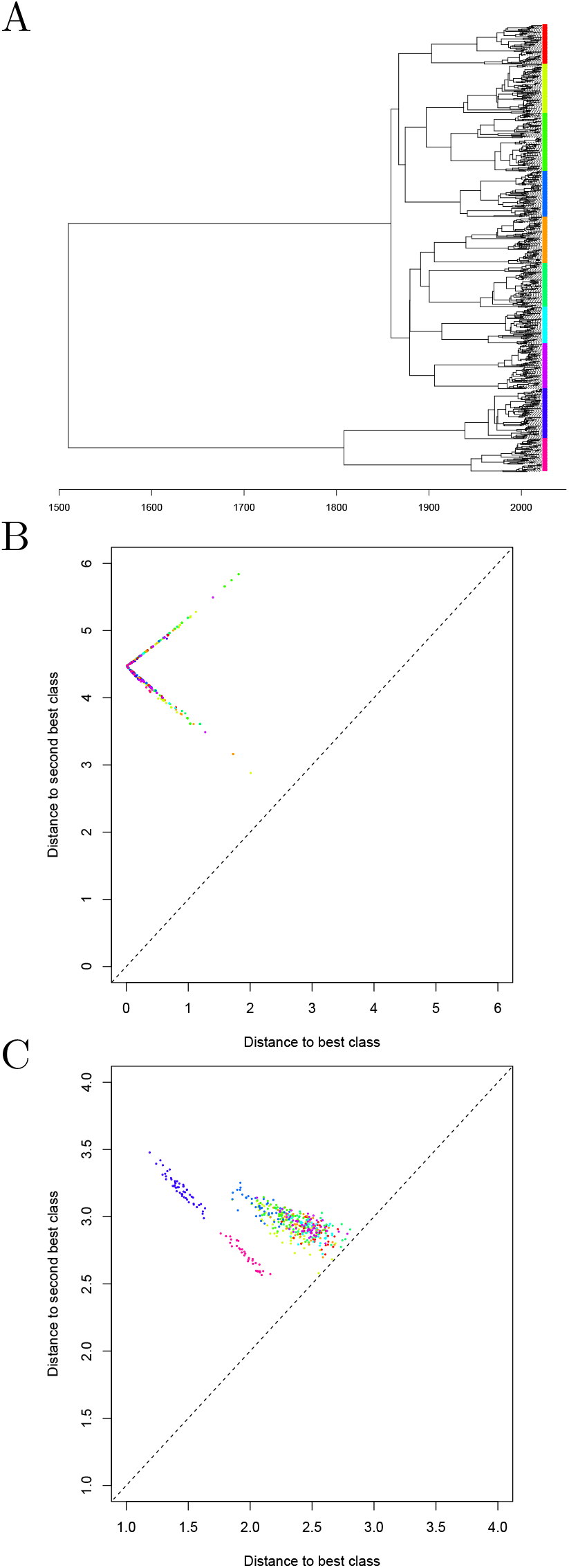
Analysis of simulated dataset of 1000 tuberculosis genomes. (A) Genealogy used for the simulation. The leaves of the tree are shown using unique colors for each of the ten lineages. (B) Results of the classifier using whole-genome sequences. Each test genome is shown as a dot, with the x-axis showing the distance to the closest class and the y-axis the distance to the second closest class. (C) Same as part (B) but based on raw read sequences rather than whole-genome sequences.

We first built the classifier illustrated in Figure 2 using the whole genome sequence data as input. Training the classifier took 3 minutes and applying it to the test genomes took 4 minutes. We computed the distance of every test genome to each of the classes. For each genome the class that had the lowest distance was the correct one, which means that all genomes were correctly classified. A measure of classification certainty can be obtained by comparing the distance from each genome to the classes with the lowest and second lowest distance. Figure 3B shows each test genome as a dot, with the x-axis representing the distance to the class with the lowest distance, and the y-axis representing the distance to the class with the second-lowest distance. In every case the distance to the closest (correct) class was much smaller than the distance to the second closest class, meaning that the classification was confident for every genome.

We then attempted to classify the same dataset based on raw read data rather than the whole genome sequences. In order to do so, simulated next-generation sequencing (NGS) data was generated for each genome using ART [40], emulating Illumina HiSeq 2500 paired-end reads of length 150bp with an average coverage of 20 fold. Training the classifier took 32 minutes and applying it to the test genomes took 17 minutes. Once again all the genomes were correctly classified. However, Figure 3C shows that for a few genomes the distance to the closest class was sometimes only slightly shorter than the distance to the second closest class. This was especially true for genomes belonging to a lineage that had a closely related lineage in the original phylogeny shown in Figure 3A. Overall, this example illustrates `KPop`’s apparent ability to successfully cluster together in abstract space sequences that share a high degree of similarity.

For comparison, we ran `mash` [28] on the simulated whole genome sequences, which took approximately 10 minutes to run. Based on the distances derived by `mash` between pairs of genomes we constructed a *k*-nearest neighbour classifier [41] with *k* = 5. We found that only 13% of genomes were correctly classified. This lack of accuracy can be traced to a lack of resolution in the result, as can be seen in the MashTree [42] output shown in Figure S1. We also applied `sourmash` [29] to the simulated raw read data, which took approximately 3 hours to run. Subsequent classification by `sourmash tax` achieved 22% accuracy at *k* = 31 (the default) and 33% at *k* = 21. While results seem to improve slightly when a smaller value of *k* is selected, inspection of confusion matrices for the classifier (Figures S2 and S3) shows a general high level of misclassification with no apparent pattern. A neighbour-joining tree based on the distances estimated by `sourmash` at *k* = 31, which has a similar quality to the one produced by `mash`, is shown in Figure S4.

### Classification of *Mycobacterium* sequencing read data

We collected a high-throughput sequencing dataset containing 1318 *Mycobacterium* genomes, all of which are available on the Sequence Read Archive [43] (Table S1). Each genome belongs to one of the 9 species *M. abscessus, M. aυium, M. boυis, M. canettii, M. caprae, M. orygis, M. smegmatis, M. tuberculosis* and *M. ulcerans*. Furthermore, *M. boυis* was subdivided into whether the genomes were derived or not from the Bacille Calmette-Guèrin (BCG) and *M. tuberculosis* was also subdivided into the 7 lineages L1, L2, L3, L4, L5, L6 and L7. There were therefore a total of 16 classes in this dataset, and the class of each genome was taken either from BIGSdb [6], TB-Profiler [44], or from previous publications [45]. The BCG genomes were manually selected and downloaded from SRA.

A data preprocessing workflow was applied to all the genomes (see Figure 4 and Methods) to filter out contaminations, i.e. reads not originating from the *Mycobacterium* genome being sequenced. Briefly, all the reads for each sample were adaptor- and quality-trimmed with TrimGalore [46] and subsequently split into chunks of 25nt. The chunks were mapped with the GEM mapper [10] version 3 onto a “pangenome” obtained by downloading from GenBank [5] all the complete genomes for the 9 species considered. Each paired-end read was kept when both ends had a length ≥ 50nt after trimming, and at least 3 chunks had been mapped to the *Mycobacterium* pangenome. The surviving reads were subsequently merged with `flash` [47] and their spectrum computed with `KPopCount` (or their sketch computed with `sourmash` for benchmarking). Samples for which less than 5*×*10^7^ *k*-mers were left after this procedure (corresponding to a coverage of roughly 10*×*) were discarded. A similar workflow was applied to samples for which single-end reads were available.

**Figure 4.**
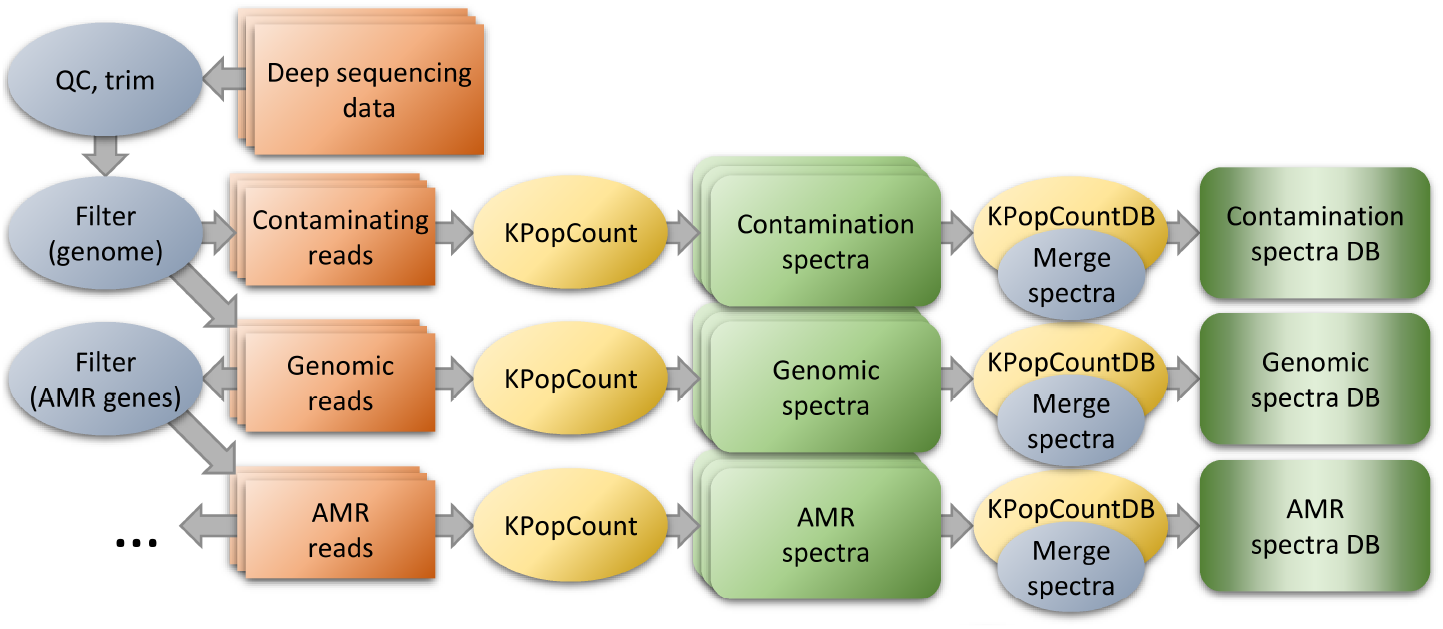
Data preprocessing workflow. When analysing NGS datasets with `KPop`, one can optionally pre-process sequencing reads in order to eliminate biases and/or have the method focus on specific parts of the genome. For instance, one might align reads to a (pan-)genome and separate them into reads that align (likely to originate from the organism being studied) and reads that do not (likely to come from contaminations). Furthermore, reads that do map to the pan-genome might be separated into groups specific to different genomic features; for instance, one might align them to a set of MLST genes or AMR genes. Full *k*-mer spectra would then be separately obtained from each group of reads (contaminations, pan-genomic, MLST genes, AMR genes) and given as input to downstream/classification methods. The choice of the group of reads from which spectra are computed determines the set of sequences seen by the method, and hence the scope of the classification.

A total of 668 genomes, roughly half in each class, was used to train the classifier, and the remaining 650 genomes were used to test its accuracy. Training and testing took approximately one hour each.

Three different classifications methods were used:

1. A method based on finding the closest class in twisted space, as described in the previous section and online [35].
2. A *k*-nearest neighbour method [41] with *k* = 5 based on finding the most represented class among the *k* training sequences which are closest to the test sequence in twisted space.
3. A random forest [33] method based on the coordinates and metadata of the training sequences in twisted space.

In all cases the twister used is the one obtained by running `KPopTwist` on class representatives, which are defined as the averaged normalised linear combinations of the *k*-mer spectra of all the sequences belonging to each class (see Methods and [35]). The results of the classification of the test set with method 1 are summarised in Table 1. For 630 of the genomes (97%) the correct class was identified as the closest one, with a further 14 genomes (2%) for which the correct class was found to be the second closest one. This left only six genomes (*<* 1%) for which the correct class was neither the first nor the second closest one. The results of the 5-NN classification (method 2) are also shown in Table 1. For 634 of the genomes (98%) the correct class was found with high probability (all nearest neighbours belonging to the correct class), and for a further eight genomes (1%) the correct class was found with lower probability (class determined by the majority rule as the nearest neighbours belong to different classes). This left only respectively five and three genomes for which an incorrect class was identified, with low and high probability (majority of the neighbours, or all the neighbours, belonging to the wrong class), respectively. Finally, Table 1 also shows the results of method 3, the random forest classifier [33], in which only eight genomes (1%) were misclassified.

**Table 1.** Results of the *Mycobacterium* classification using distance to class, using *k*-nearest neighbours and using a random forest.

| Lineage | Number | Distance by class | | | $k$-nearest neighbours | | | | Random forest | |
| --- | --- | --- | --- | --- | --- | --- | --- | --- | --- | --- |
|  |  | First correct | Second correct | Neither | Correct, certain | Correct, uncertain | Incorrect, uncertain | Incorrect, certain | Correct | Incorrect |
| *M. abscessus* | 49 | 49 | 0 | 0 | 49 | 0 | 0 | 0 | 49 | 0 |
| *M. avium* | 34 | 34 | 0 | 0 | 32 | 2 | 0 | 0 | 34 | 0 |
| *M. bovis* | 50 | 44 | 4 | 2 | 50 | 0 | 0 | 0 | 49 | 1 |
| *M. bovis* BCG | 47 | 47 | 0 | 0 | 46 | 0 | 1 | 0 | 47 | 0 |
| *M. canettii* | 5 | 4 | 0 | 1 | 4 | 0 | 0 | 1 | 3 | 2 |
| *M. caprae* | 50 | 44 | 5 | 1 | 49 | 1 | 0 | 0 | 50 | 0 |
| *M. orygis* | 8 | 8 | 0 | 0 | 8 | 0 | 0 | 0 | 7 | 1 |
| *M. smegmatis* | 21 | 21 | 0 | 0 | 21 | 0 | 0 | 0 | 21 | 0 |
| *M. tuberculosis* L1 | 55 | 54 | 1 | 0 | 51 | 4 | 0 | 0 | 55 | 0 |
| *M. tuberculosis* L2 | 50 | 47 | 2 | 1 | 47 | 0 | 2 | 1 | 49 | 1 |
| *M. tuberculosis* L3 | 52 | 51 | 1 | 0 | 51 | 0 | 0 | 0 | 51 | 1 |
| *M. tuberculosis* L4 | 51 | 49 | 1 | 1 | 48 | 0 | 2 | 1 | 49 | 2 |
| *M. tuberculosis* L5 | 51 | 51 | 0 | 0 | 51 | 0 | 0 | 0 | 51 | 0 |
| *M. tuberculosis* L6 | 50 | 50 | 0 | 0 | 50 | 0 | 0 | 0 | 50 | 0 |
| *M. tuberculosis* L7 | 28 | 28 | 0 | 0 | 28 | 0 | 0 | 0 | 28 | 0 |
| *M. ulcerans* | 49 | 49 | 0 | 0 | 49 | 0 | 0 | 0 | 49 | 0 |
| Total | 650 | 630 | 14 | 6 | 634 | 8 | 5 | 3 | 642 | 8 |

The three genomes that were misclassified with high probability in the *k*-nearest neighbours analysis were among the six genomes for which the correct class was neither the first or second closest one, and among the eight genomes misclassified according to the random forest method. According to the original metadata for our dataset, these three genomes belonged to *M. canettii, M. tuberculosis* L2 and *M. tuberculosis* L4, respectively. However, there might have been inaccuracies in these classifications, as suggested by the fact that some of the designations in BIGSdb [6] did not always agree with the results of TB-Profiler [44]. Such inaccuracies would make our results look worse than they are if they are present in the test set, but, more importantly, they could affect classification result even for test genomes that are correctly labeled if they are present in the training set. Overall, it is likely that a better curated dataset would produce an even more accurate classifier. The number of reads available for the different samples spans almost three orders of magnitude, but this did not affect the results, which shows that `KPop` works in a way which is largely independent of the sequencing technology and coverage level.

We also processed the same dataset using `sourmash tax` [29]. The overall accuracy of the classification was 31% at both *k* = 31 and *k* = 21, with results being virtually independent of the choice of *k*. Inspection of the confusion matrices (Figures S5 and S6) shows that this classifier worked well for some *Mycobacterium* species but was especially inaccurate at distinguishing between the different lineages of *M. tuberculosis* ; in particular, `sourmash tax` seems to mistake most of the tuberculosis genomes for *M. boυis* BCG. Given that the misclassified sequences are very similar, this result suggests once again the lack of fine resolution by MinHash-based methods as the most likely reason for the poor performance of `sourmash` on this dataset.

### Classification of simulated genomes from a recombinant population

Unlike typical bacterial pathogens, *Mycobacterium tuberculosis* does not recombine and does not have much genome content variability [48]. We simulated a dataset of 100 genomes from a population in which the recombination/mutation ratio is equal to one and in which gene gain and loss is frequent, similar to bacterial species such as *Salmonella enterica, Escherichia coli* or *Clostridioides difficile* [48]. An ancestral recombination graph was simulated using SimBac [49], which included the clonal genealogy shown in Figure S3. Gene content variation was added using SimPan [50]. Finally, NGS data was generated for each genome using ART [40] as in the previous simulated example. There were three clear clusters in the simulated dataset (Figure S7), and we used half of the genomes from each cluster as a training set and the remainder as a test set.

Both `KPop` and `sourmash tax` [29] achieved perfect classification in this case, showing how *k*-mer based methods are largely independent of the effects of recombination in the population.

In order to shed light on the quality of the clustering generated by the two methods, we again investigated how many of the five nearest neighbours identified by `KPop` and `sourmash` in the training set belong to the same class as that of each test sequence. The results are shown in Table 2. While all the nearest neighbours identified by `KPop` always belong to the correct simulated lineage, that is sometimes not the case for `sourmash` at *k* = 31, pointing to a better quality of the distances produced by `KPop` in this scenario.

**Table 2.** Distribution of the test sequences by number of nearest neighbours being in the same class as that of the test sequence, as found by KPop and sourmash [29] in the simulated recombination example.

| Lineage | Number | NNs in the same class |  |  |  |
| --- | --- | --- | --- | --- | --- |
|  |  | 5 | 4 or 3 | 2 or 1 | 0 |
| KPop |  |  |  |  |  |
| 1 | 25 | 25 | 0 | 0 | 0 |
| 2 | 11 | 11 | 0 | 0 | 0 |
| 3 | 13 | 13 | 0 | 0 | 0 |
| Total | 49 | 49 | 0 | 0 | 0 |
| sourmash |  |  |  |  |  |
| 1 | 25 | 19 | 6 | 0 | 0 |
| 2 | 11 | 3 | 4 | 4 | 0 |
| 3 | 13 | 6 | 3 | 3 | 1 |
| Total | 49 | 28 | 13 | 7 | 1 |

### Classification of simulated SARS-CoV-2 whole genome sequences

We simulated a dataset of 10,000 whole genome sequences of SARS-CoV-2. First, a genealogy was simulated with sampling happening throughout year 2021, from 100 lineages that emerged in the second half of year 2020 and shared a last common ancestor in early 2020 (cf Figure 5A). Since the lineages emerged at different times, some were sampled more than others, with the number of samples for each lineage ranging between 2 and 309. Genomes of length 29,903 bp were generated by mutation on the branches of this tree, using as starting point the reference genome Wuhan-Hu-1 [51] and applying a substitution rate of 10^*−*3^ per year per site [52].

**Figure 5.**
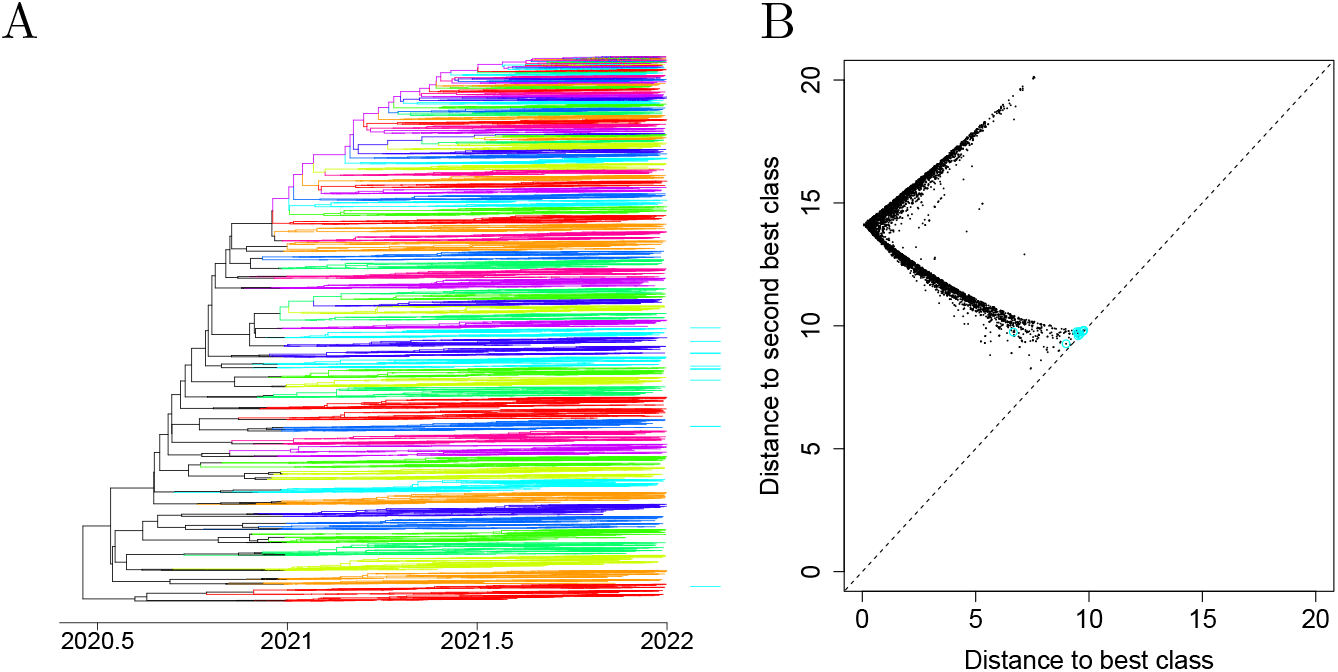
Analysis of simulated dataset of 10,000 SARS-CoV-2 genomes. (A) Genealogy used for the simulation. The branches of the tree are colored non-uniquely according to lineages. (B) Results of the classifier. Each test genome is shown as a dot, with the x-axis showing the distance to the closest class and the y-axis the distance to the second closest class. The 11 genomes for which the closest class was not the correct class are highlighted in cyan on both parts.

Within each lineage, a randomly selected half of the genomes was used to train the `KPop` classifier and the remaining half to test it. Training the classifier took about 8 minutes and applying it to the test set took about 4 minutes. Figure 5B shows for each of the test genomes the distance to the closest and second closest classes. For the vast majority (99.8%) of the genomes, the closest class corresponded to the expected one and hence the classification was correct. There were only 11 genomes for which this was not the case, highlighted in cyan in Figure 5B. Many of these had similar distances between the closest and second closest classes, with the latter being the correct class. The only genome for which that was not the case was from a small class with just three representatives, two of which were used for training and one for testing, with the test genome happening to be more dissimilar to the two training genomes than to other genomes of other classes.

### Classification of SARS-CoV-2 sequences

We also reanalysed the Pangolin dataset of COVID-19 lineage designations [53, 54]. As of the time of our analysis, this global dataset contained 1,285,005 full genome sequences of SARS-CoV-2 classified into 1636 lineages. The number of sequences per lineage was very unbalanced, ranging from 3 (for lineage C.6) to 68,299 (for lineage AY.122). We used half of the sequences in each lineage to train the classifier and the other half for testing. Training and testing took about two hours each on a 64-core HPC node with hardware specs similar to the ones previously mentioned. 93.6% of the test sequences turned out to be closest to the correct class (Figure 6); in addition, the correct class was the second closest class for 2.9% more sequences. Furthermore, if we consider only the sequences for which the smallest distance is at least 10% smaller than the second smallest distance, we find that 87.3% of the test sequences meet this criterion and most (98%) of these sequences have the smallest distance to the correct lineage. In other words, as expected, the accuracy of the classification is higher for lineages for which the classifier is more confident (Figure 6).

**Figure 6.**
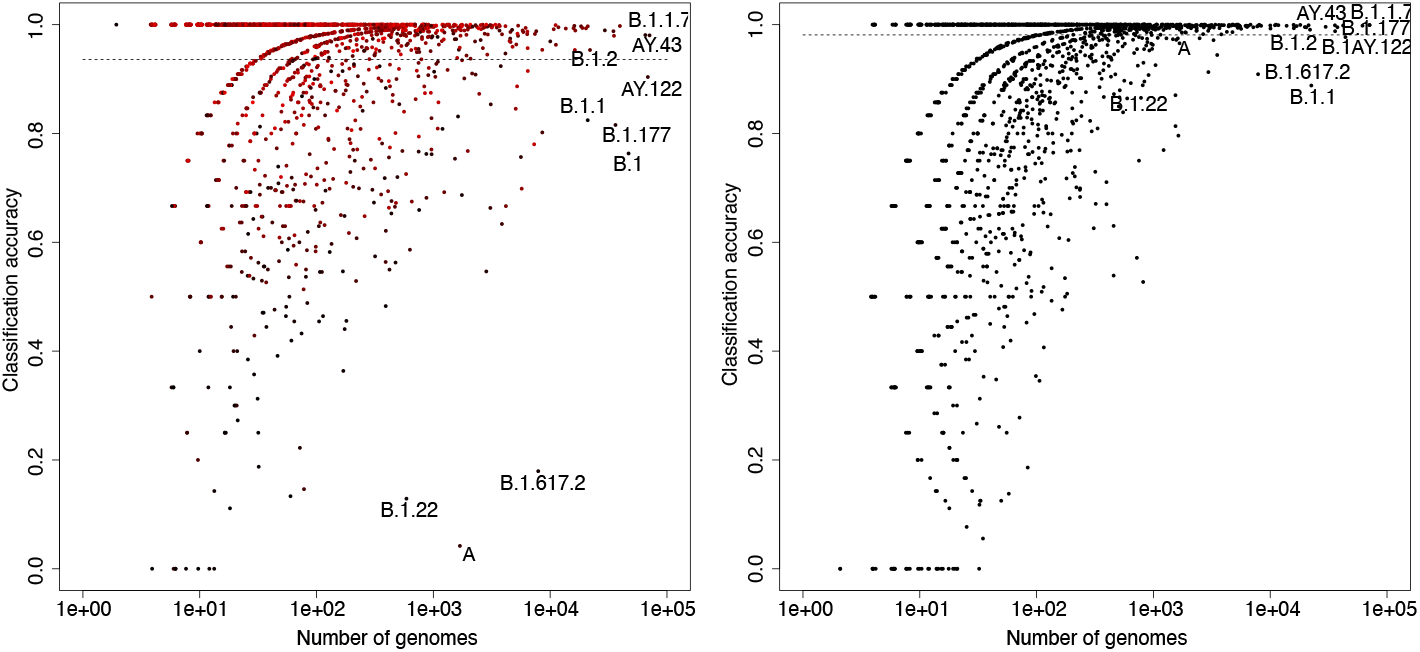
Analysis of Pangolin SARS-CoV-2 dataset. Each point represents a lineage; the x-axis shows the number of genomes, and the y-axis the proportion of correctly classified genomes. The left panel shows the result of classifying according to distance to class; the right panel shows the result of classifying according to a random forest. On the left, the colours represent the level of confidence in the classification, ranging from black to red proportionally with the difference in distance between the first and second closest classes. Black therefore represents low confidence and red high confidence in the classification.

For comparison, we applied Usher [55] following the instructions at https://usher-wiki.readthedocs.io/en/latest/UShER.html to the same set of genomes, using the same subsets for training and testing. Using a 5-nearest neighbour classifier based on the phylogeny produced by Usher, we found that 97.5% of the test sequences were correctly classified. However, Usher needed more than 100 times more computational resources than `KPop`. On top of that, Usher used as input a phylogeny of the training sequences with 600,000 leaves which would have needed even more time to be generated.

We note that cases of misclassification are often explained by heterogeneity whereby a child class is not simply defined, phylogenetically speaking, as a subtree of its parent. This is illustrated in Figure S8 for lineage B.1.617.2 and its child lineage AY.4 (alias of B.1.617.2.4). In this case, we see that AY.4 does not simply correspond to a sub-branch of B.1.617.2; instead, it is defined as the union of several separate sub-branches. This translates into the training clusters not being hyperspherical in twisted space, which makes classification based on class centroids non-optimal. We therefore tried `KPop` classification based on random forests [33] and found that a naive classifier (with *n* = 240 trees) had an accuracy of 98.1%. For this paper we did not consider other possibilities, even though more sophisticated methods, such as support vector machines or AI algorithms, might perform even better.

### Identification of closely related SARS-CoV-2 sequences

Many of the workflows described in the previous sections are based on the computation of distances between twisted *k*-mer spectra, once a twister suitable for classification has been created. This observation naturally suggests the possibility of using our framework to store a large number of twisted sequences into a database and, once the twisted spectrum for a new sample becomes available, identify the sequences in the database that are closest to the new one. As explained in the Methods, this capability is natively provided by `KPopTwistDB`; for instance, it lies at the heart of the *k*-NN based classifier for *M. tuberculosis* presented above. After identification of nearest neighbours, the new twisted spectra can be added to the database (see Figure 7 for a complete description of the workflow).

**Figure 7.**
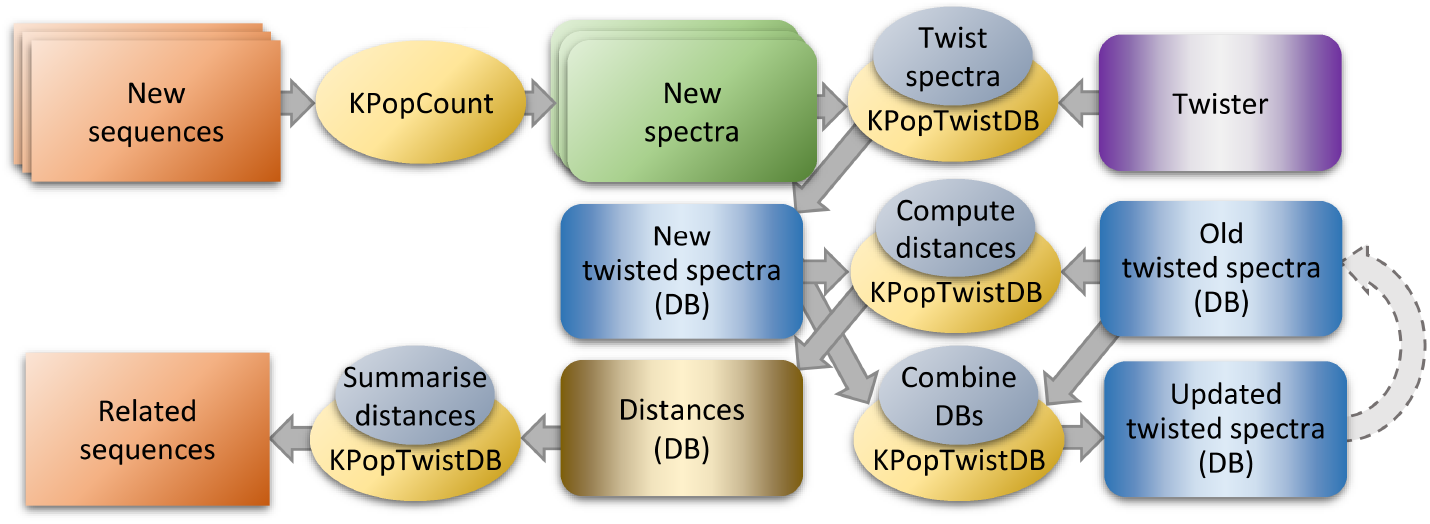
Relatedness engine workflow. The workflow is very similar to the classification one depicted in Figure 2, and can be considered a derivative of it — given a classification workflow, one can always automatically obtain a workflow determining relatedness according to the criteria used to classify sequences. After twisting new sequences according to the classifying twister found according to the procedure of Figure 2, one uses `KPopTwistDB` to compute the distances between them and a set of twisted spectra previously stored in a database. Such distances can be summarised with `KPopTwistDB` in order to identify a set number of closest sequences in twisted space. One would then use `KPopTwistDB` a third time in order to add the new twisted spectra just obtained to the database of old twisted spectra.

Using the same COVID-19 dataset and twister described in the previous section, we tested this idea by finding the 300 closest neighbours of a number of test sequences in a binary `KPop` dataset containing the roughly 600,000 sequences used to train our classifier. The database was 8.4 GB in size — we did not test the possibility of splitting the sequences into smaller files, which might help to achieve a better parallelism. The database took about 2 minutes to load into memory (which, like for the index in in BLAST or other alignment programs, only happens once at the beginning of the workflow, independent of the number of sequences in the query). After that, determining the 300 closest neighbours took about 20 seconds for each sequence. The results were consistent with expectations, with most of the closest sequences identified belonging to the same class as that of the query.

### Taxonomic classification

Finally, we devised a modified `KPop`-based classification workflow that uses taxonomic information to attribute each genome to all taxonomic levels it belongs to. One separate classifier is generated and used for each taxonomic level, and test sequences are attributed to the lowest level that is consistent with predictions generated by all other higher-level classifiers. When the correct class is missing, for example a species present in the test dataset that was not found in the training dataset, the classifier returns the closest match at the lowest possible taxonomic level.

Data from the TARA Oceans metagenome project [56] was used by a recent study [57] to create a non-redundant database of archaeal, bacterial and eukaryotic genomes. This database includes 957 metagenome-assembled genomes, each of which having partial taxonomic classification. 479 genomes were used for training and the remaining 478 were used for testing. We applied both our `KPop` workflow and `sourmash tax` to the TARA dataset, and the results are shown in Table 3. The `sourmash` error rate was very low, but this came at the cost of very conservative results, in which even at at the domain level (i.e., bacteria, eukaryota, or archaea) most genomes were left unclassified. The classification obtained using `KPop` was more assertive, with, for instance, more than 90% of genomes classified at the domain level. The percentage of uncalled increased for both algorithms as finer taxonomic classification was explored; that is expected, given that at lower taxonomic levels there were less and less repeated classes within the training and test sets.

**Table 3.** Results of the taxonomic classification using sourmash and KPop for the TARA Oceans database.

| Classification using **KPop** |  |  |  |  |  |  |  |
| --- | --- | --- | --- | --- | --- | --- | --- |
|  | Domain | Phylum | Class | Order | Family | Genus | Species |
| Total | 478 | 438 | 367 | 267 | 148 | 54 | 7 |
| Correct | 435 | 225 | 143 | 96 | 59 | 18 | 3 |
| Incorrect | 6 | 18 | 13 | 8 | 1 | 0 | 0 |
| Uncalled | 37 | 195 | 211 | 163 | 88 | 36 | 4 |
| Classification using sourmash tax |  |  |  |  |  |  |  |
|  | Domain | Phylum | Class | Order | Family | Genus | Species |
| Total | 478 | 438 | 367 | 267 | 148 | 54 | 7 |
| Correct | 95 | 61 | 54 | 42 | 31 | 16 | 0 |
| Incorrect | 2 | 0 | 0 | 0 | 0 | 0 | 0 |
| Uncalled | 381 | 377 | 313 | 225 | 117 | 38 | 7 |

## Discussion

We have introduced `KPop`, a novel methodology for the comparative analysis of large datasets of microbial genomes. `KPop` does not require a sequenced sample to be assembled or aligned to a reference genome, and uses the full spectra of *k*-mers found in a set of sequences or raw reads. By applying to this highly dimensional *k*-mer space an appropriate, dataset-specific transformation (the “twister”) based on Correspondence Analysis, it becomes possible to compare very large numbers of genomes in a twisted space of reduced dimension. For example, one can classify genomes into groups or find close relatives at scale and with high precision.

In particular, `KPop` lends itself well to highlighting differences at the sub-species level even when the overall genomic diversity is low, as demonstrated by our exploration of a number of scenarios (simulated recombination of close species, large simulated and real datasets for *M. tuberculosis* and SARS-CoV-2). This ability shown by `KPop` should be compared with MinHash-based methods such as `mash` [28] or `sourmash` [29], which are arguably the most popular methods relying on *k*-mers currently in use. Although from a purely computational standpoint they are even more scalable than `KPop` thanks to their use of small signatures (“sketches”), they are also limited in their accuracy, for example when classifying genomes into lineages within species [28]. In contrast, `KPop` takes as its starting point full spectra of (shorter) *k*-mers; as the results we present here demonstrate, this choice makes it natural for us to directly operate on short unassembled reads, gives excellent accuracy, retains enough scalability for most applications in genomic microbiology, and produces classifiers that outperform `mash` [28] and `sourmash` in all the scenarios we tested. In fact, our results clearly point out to our `KPop` being better at evaluating genomic distances than MinHash-based methods, in particular when small differences need to be resolved, and suggest that `mash` [28] or `sourmash` should be used with extreme caution, or not used at all, when high-resolution results are needed. It should be noted that the research on MinHash-based methods is an active field, and many alternative ways of computing sketches and/or distances out of them have been proposed [58, 31, 59, 60].

`KPop` offers another even more important conceptual advantage. While one can compare it with other methods thanks to its ability to evaluate genomic distances, `KPop` is natively capable of embedding *k*-mer spectra in a reduced-dimensionality space. This makes it possible to transform each genome into a vector, and feed such vectors straight away to classifiers based on machine-learning or AI approaches. In particular, one could readily leverage existing high-performance software developed in the AI domain by supplying the embeddings produced by `KPop` to vector databases such as Milvus [61] and by implementing relatedness engines for large collections of genomic data on top of similarity search libraries such as Faiss [62].

Compared to other methods that are specifically designed to perform a single task such as establishing relations [28] or searching for specific genetic elements [24], our strategy retains superior flexibility. Thanks to its implementation as a set of modular programs that can easily be combined into a number of different workflows, one can explore a number of scenarios. In particular, more complex workflows than the ones presented here can be easily performed and automated in our framework, without sacrificing scalability — all computer-intensive steps are naturally parallelised whenever possible, so that very large datasets can be analysed within a short wall-clock time frame given enough CPUs and memory.

The fact that `KPop` can model relations between genomes efficiently and accurately, and more accurately than minimizers-based methods, makes it a natural starting point for future methodological studies to try and develop scalable and high-precision methods for microbial phylogenetics [63]. We intend to explore these ideas in future work.

## Methods

### General strategy

The general philosophy of `KPop` is summarised in the first section of the Results. A more detailed graphical description of `KPop` classification in twisted space can be found in the left-hand part of Figure 8.

**Figure 8.**
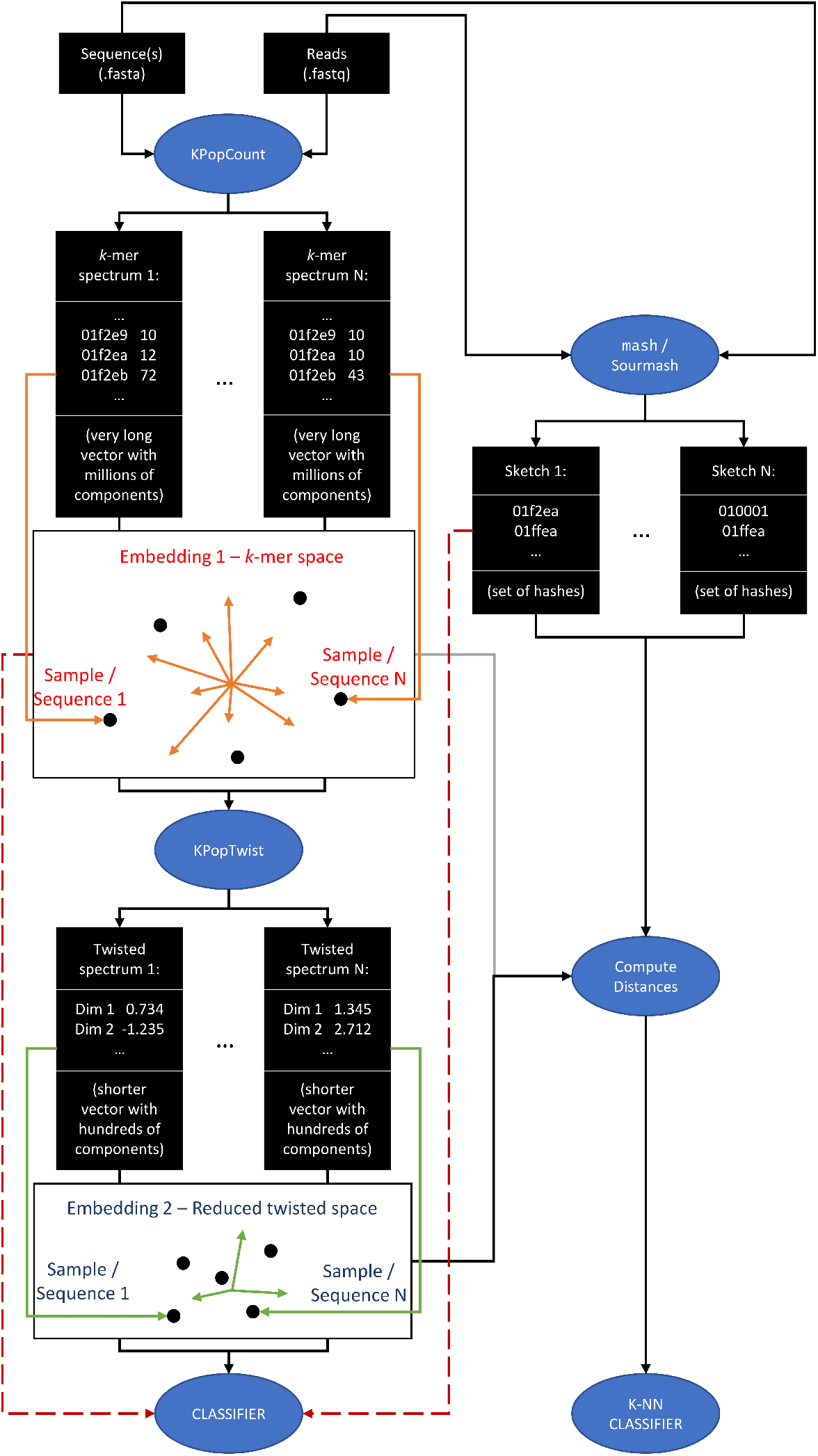
General KPop classification strategy. First, KPopCount computes *k*-mer spectra, which for the typical values used by KPop are very long vectors with millions of components. Second, `KPopTwist` “twists” such vectors to Correspondence Analysis (CA) space, which produces much shorter vectors (of the order of the number of samples or sequence classes/clusters - the effective number might be even smaller because of the negligible inertia they carry in CA sense). While distances can be computed both from spectra and twisted vectors (grey and black line on the left of “Compute Distances”), feeding spectra as vectors directly to classifiers would be problematic due to the their high dimensionality (red arrow on the left). However, twisted vectors can be used as direct input to classifiers thanks to the CA-based dimensionality reduction step.

### Implementation and availability

For the abstract methods described above to be turned into practice, a number of practical steps need to be performed. We have implemented them as a complete suite of modular programs. They are mostly written in OCaml [64] and partly as R [65] scripts, and can be freely downloaded as source code from [35]. Compilation from scratch and installation are easy; alternatively, pre-compiled binaries for some platforms are available from [35] or from `bioconda` (package `KPop`). The programs performing heavy-duty computation (mainly `KPopCountDB`, `KPopTwist`, and `KPopTwistDB`) have built-in parallelisation support to reduce the overall wall-clock execution time.

### KPop programs

The workflows used in this paper rely upon four main programs in the `KPop` suite:

#### KPopCount

It allows to extract *k*-mer spectra from FASTA and single- or paired-end FASTQ files. Spectra are output in textual format

#### KPopCountDB

It allows to collect *k*-mer spectra into binary databases, and export the resulting tables as either binary or text files. A number of transformations can be performed on the content of the spectral database

#### KPopTwist

Given a database of *k*-mer spectra, it allows to generate a coordinate transformation based on Correspondence Analysis [66, 67] that implements an unsupervised dimensional reduction optimised for that database. Such transformation (the “twister” in `KPop` jargon) turns (“twists”) a *k*-mer spectrum into a typically much smaller numerical vector; the transformation can be stored in binary format and applied to other spectra in the future

#### KPopTwistDB

It is the Swiss knife of twisted spectra. Among other things, it can be used to twist *k*-mer spectra if a suitable twister is provided; to collect twisted spectra into databases that can be output as either binary or text files; to compute and summarise distances between twisted spectra.

Figure 9 summarises the main transformations between data types and file formats performed by the four programs. More details about the methods used in each program follow.

**Figure 9.**
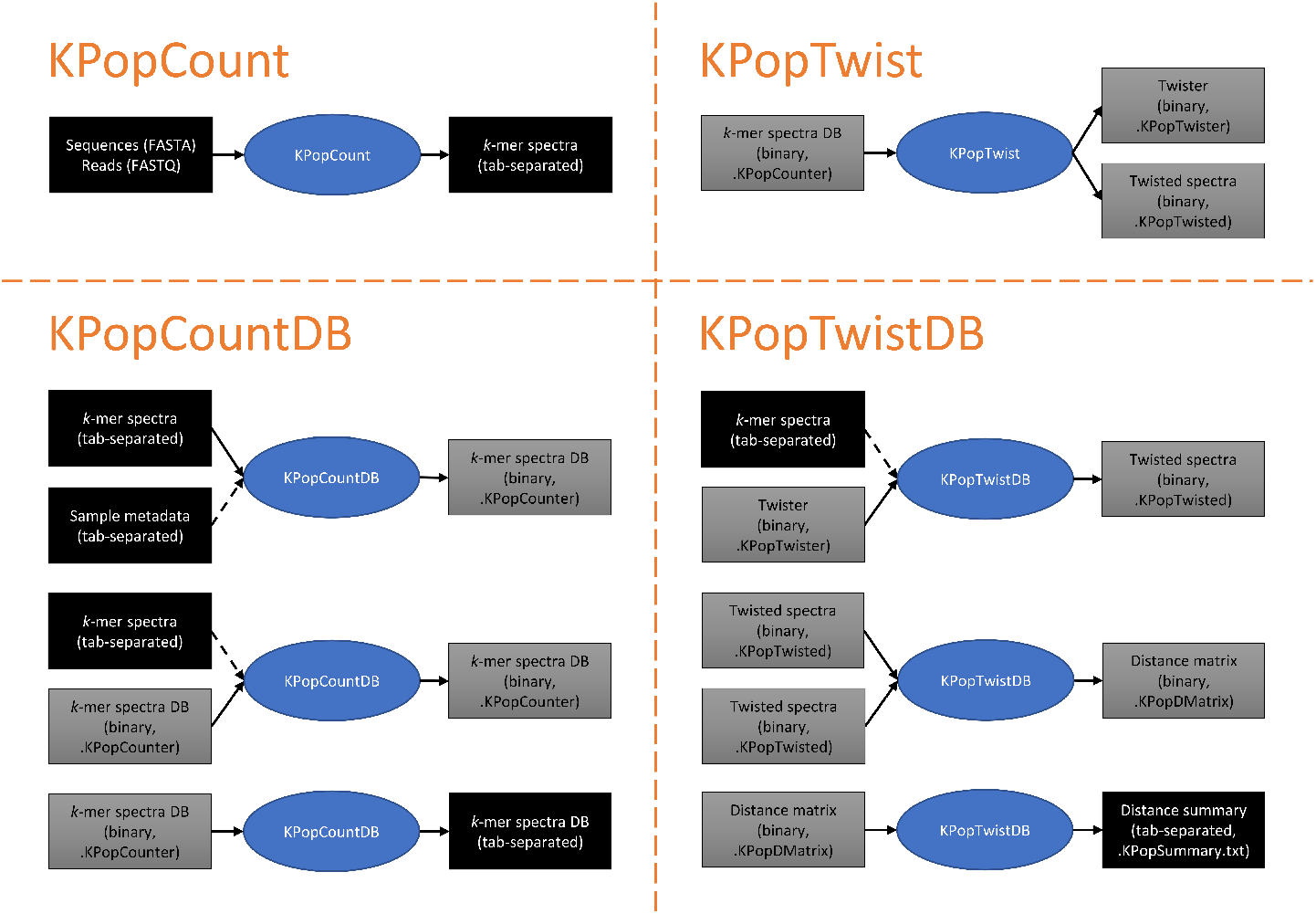
Main data transformations performed by the KPop programs. In the case of `KPopCountDB` and `KPopTwistDB`, the list is not exhaustive. For instance, with `KPopCountDB` it is possible to compute distances between untwisted *k*-mer spectra; with `KPopTwistDB`, one can accumulate twisted *k*-mer spectra or distances into existing databases, and convert binary files from/to plain-text tab-separated tables. In the spirit of BLAST makeblastdb, automatic naming rules are applied to files, and one only needs to supply file prefixes.

### KPopCount

`KPopCount` accepts as input a number of single- or paired-end files FASTA/FASTQ files, and counts the *k*-mers of a specified size present in them. At the moment regular *k*-mers are used, although support for other schemes (notably gapped *k*-mers [68]) might be explored in the future. `KPopCount` accepts both DNA and protein sequence. To cope with the lack of directionality of most sequencing protocols, the reverse complement of the input sequences is automatically generated and taken into account when counts are computed for DNA. DNA spectra are subsequently de-duplicated by computing the reverse complement of each *k*-mer and only keeping the lexicographically smaller of the two. The counts for palindromic *k*-mers are halved.

#### Choice of k

It should be noted that the choice of the correct value for *k* depends on the problem at hand and is not entirely straightforward. In particular, it is recommended that saturation analysis be conducted while the analysis workflow is set up; i.e., *k* should be chosen as the smallest values for which the results become stable. That is because when using too big a *k* most *k*-mers will be either absent or present with frequency 1, resulting in each of them having the same relevance and making the spectrum sensitive to noise, thus ultimately leading to overfitting; while, when using a value of *k* which is too small for the degrees of freedom of the system being studied, the generated *k*-mer spectrum will not have enough resolution to produce sufficiently sharp frequency peaks that the CA step can focus on to identify relevant features. As a rule of thumb, choosing

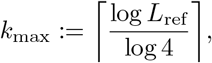

where *L*_ref_ is the size of the reference genome, should provide an upper bound for *k*, as such a choice guarantees that for a random genome most *k*-mers would be present with low frequency; in practice, the optimal value of *k* might differ slightly from *k*_max_. Note that in most cases one does not need to possess an explicit assembly of the sample to be able to estimate *L*_ref_; typically, *L*_ref_ will be of the order of the size of the genome being studied.

A more general approach that does not rely on prior knowledge and might provide a stricter bound can be obtained by selecting a value of *k* that maximises the condition number of the spectra and hence also the discriminating power of the method. In practice, as the minimum *k*-mer frequency is often 1 — in particular when noisy raw sequencing reads are considered — one might instead maximise the ratio between maximum *k*-mer frequency and average coverage across all samples. I.e., if 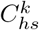 is a non-zero count for *k*-mer *h* in sample *s* and there are *n*_*h*_ non-zero k-mers, one might choose

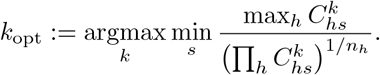

This criterion would also work in the general case of metagenomic samples for which the size of the underlying reference is often unknown.

### KPopCountDB

`KPopCountDB` can be used to perform operations on collections of *k*-mer spectra.

#### Table structure

Spectra are stored as columns of a “database”; conceptually, a database can be seen as the binary representation of a matrix, although additional data structures are present in practice to make the matrix extendable and searchable. New columns in the format produced by `KPopCount`, or new metadata in tabular format, can be added to an existing matrix.

In more detail, the general conceptual layout of a database is as follows (we drop the implicit dependency of counts *C* on *k* in the notation):

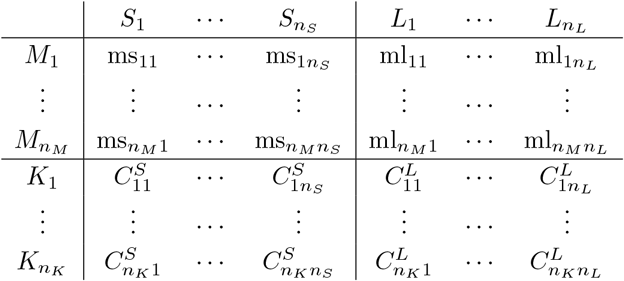

In this example, 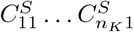 would be the *k*-mer spectrum corresponding to sample 1; it would have *n*_*K*_ *k*-mers.

A `KPop` *k*-mer database can also be augmented with additional information. In particular, linear combinations of existing columns/data points can be generated and stored as additional columns of the matrix — one such combination being the vector 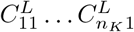 in the example above. This feature allows, for instance, to generate and store an average representative spectrum for a set/class of sequences as the linear combination of the spectra of all the sequences belonging to that class. Such linear combinations can then be used as active or passive points in subsequent analyses. Additionally, metadata (typically as text strings, labelled 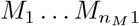 in the figure) can be stored as additional rows for each sample, or added at a subsequent time.

The table can also be output as text. All expensive operations, including output, are natively parallelised in order to reduce the overall wall-clock time spent in them.

#### Transforming frequencies

According to common experience in the field of *k*-mer statistics, the *k*-mer distribution follows a curve whereby there is a large number of low-frequency *k*-mers, typically due to sequencing noise. `KPopCountDB` supports clean-up of the spectra by implementing a number of column-based transformations of the (*C*^*S*^)(*C*^*L*^) lower part of the database.

Different from what happens in detection-oriented methods, our approach does not primarily attempt to reduce the complexity of signatures – while using longer *k*-mers might allow to find sequences that are distinctive of, and specific to, some particular genome, in our case the information is captured in the overall vector of *k*-mer frequencies. However, it is desirable to eliminate the noise given by low-coverage *k*-mers, which is bound to manifest itself in any case due to the finite number of reads used to probe the sample. Depending on the genome, another problem might be given by high-frequency *k*-mers due to very repetitive regions, which might unduly capture the attention of the linear methods for dimensional reduction used downstream.

In order to tackle the latter, we use a strategy based on power-law or logarithmic mapping of the original *k*-mer frequencies. As the methods implemented in `KPopTwist` critically rely upon input data being positive, we define transformations as if we were re-mapping the original counts to new integers. Aside from possible overflow problems depending on the actual exponent of the power law, this strategy keeps the transformed vector compatible with the overall framework.

To accomplish that, if *m* is the maximum value of the vector of counts *υ* and we are applying a logarithmic transformation, we renormalise the vector by a factor *r* defined as

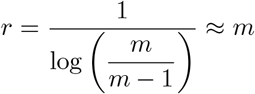

which ensures that

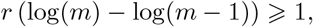

i.e. that *m* and *m −* 1 are represented by different integer numbers. Hence, and imposing that renormalised counts be 0 for frequencies equal to or less than some cutoff *c*, we redefine *υ* as

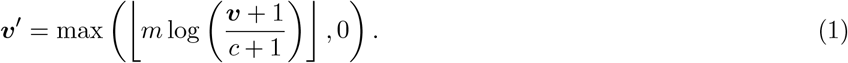

It should be noted that apart from a column-dependent proportionality factor, this transformation coincides with the centered log-ratio (*clr*) transformation that is widely used for compositional data and “binning” programs, e.g. [26]. However, different from *clr*, Equation (1) is always positive and does not suffer from the problem of zero counts. Following the same criterion, given a 0 ⩽ *p <* 1 we can define a power-law transformation

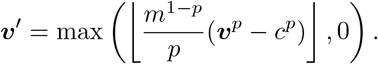

having the same property that *υ*^*l*^(*m*) *− υ*^*l*^(*m −* 1) ⩾1.

The transformations introduced so far tend to decrease the distance between samples. It is also possible to transform the data in order to *increase* the distance between samples and specificity — to accomplish that, one has to increase the relative importance of the most frequent *k*-mers. An obvious way of doing this is by performing a power-law transformation with *p* ⩾ 1. In this case, however, the representability condition becomes important for small rather than large numbers and we have to make sure that *υ*^*l*^(*c* + 1) = 1, resulting in the transformation

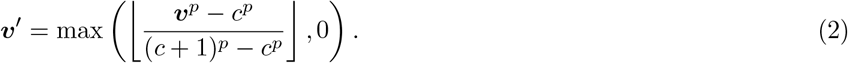

Depending on the dataset, the last transformation may incur overflow problems when values of *p* are too large. However, in our implementation we always subsequently convert to floating point and normalise the pseudocounts produced with Equation (2), which alleviates the problem.

Further cleanup would be possible at this stage, for instance by eliminating uninformative *k*-mers whose fractional frequencies are essentially constant across all samples. However, in general row-based transformation such as this one produce results that depend on the order whereby columns are added to the database, and hence we do support a few of them (mainly the removal of *k*-mer rows such that all the frequencies are below a given threshold) but only when the table is exported as text.

### KPopTwist

`KPopTwist` performs unsupervised dimensional reduction of a set of *k*-mer spectra and determines dataset-specific ways of transforming them (called “twists” in `KPop` terminology) that can be stored for use on samples obtained at a later time.

`KPopTwist` uses Correspondence Analysis (CA, [66, 67]) as the underlying procedure to identify, define and build a set of dimensions being relevant to the system being considered. CA is a long-standing statistical procedure, and it has been employed to characterise a number of systems in diverse fields ranging from econometrics to biology. Interestingly, when prototyping our method on a wide range of possible use cases we were able to determine that CA seems to consistently outperform other methods for dimensional reduction — PCA (principal component analysis) in particular, but also a number of more recent non-linear techniques. One reason might be due to *k*-mer spectra being compositional and CA a technique that is well-suited to compositional analysis [69]; another might be the fact that CA has known strong connections with network/clustering analysis [70]. One good property of CA is that, as PCA does, it ranks the resulting dimensions in order of decreasing quantitative importance; in addition, CA is an explainable procedure, in that one can trace back the definition of each relevant dimension to a definition in terms of a specific set of *k*-mers appearing in the original spectra. Taken together, these two factors would allow the identification of the *k*-mers that characterise each cluster or class highlighted by the procedure, even though we do not explore this possibility here.

Finally, `KPopTwist` also implements a random downsampling of the set of *k*-mers appearing in the spectra. That is useful to validate the robustness of dimensional reduction with a procedure reminiscent of bootstrapping.

### KPopTwistDB

`KPopTwistDB` performs a number of operations on twisted vectors. By using it one can:

- Twist *k*-mer spectra. That requires as input a sequence of spectra as produced by `KPopCount` and a twister transformation as produced by `KPopTwist`, and returns an object of type “twisted” that contains a set of vectors in twisted space
- Compute pairwise distances between two sets of twisted *k*-mers, and summarise them
- Accumulate twisted vectors or distances into larger databases. This is useful, for instance, to parallelise the creation and processing of datasets containing a large number of samples
- Convert objects of any kind (twisters, twisted, distance) from binary to tabular text form, and vice versa. The tabular representation can then be read into `R` or other programming languages for further downstream analysis.

#### Distance in twisted space and metric

Given two twisted vectors *υ* ≡ *υ*_1_ … *υ*_*d*_ and ***w***≡ *w*_1_ … *w*_*d*_, *d* being the number of dimensions of the twisted space, `KPopTwistDB` implements a generalised Minkowski distance *D*_*p*_ between them defined as

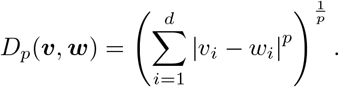

CA embeds *k*-mer spectra into lower-dimensionality spaces. Similar to what happens with PCA, such dimensions are sorted by decreasing “inertia” (a quantity analogous to variance), that we will note as ***I*** ≡ *I*_1_ … *I*_*d*_. The inertia vector produced by CA is positive and normalised, i.e., it holds true that 0 ≤ *I*_*i*_ ≤ 1 ∀*i* ∈ [1 … *d*] and 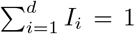 and the components of the vector are sorted in non-increasing order, i.e., it holds true that *I*_*i*_ ≥ *I*_*i*+1_ ∀*i* ∈ [1 … *d* − 1]. When computing distances between twisted vectors, it then makes sense to define a procedure that takes into account the relevance of each dimension in terms of explained inertia — i.e., we might wish the dimensions to contribute to the overall distance in a way which is somehow proportional to the amount of inertia they carry, in particular when the effective number of degrees of freedom of the system turns out to be smaller than the number of CA dimensions. In order to do so, we introduce the concept of *metric*, i.e. a vector ***µ*** that is a (positive and separable) function of ***I***; we would then compute the distance between two twisted vectors *υ* and *w* as

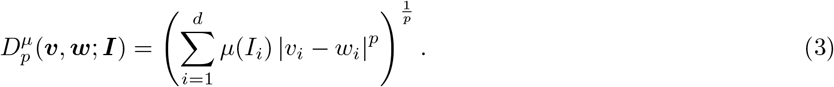

Many choices would be possible for the function *µ*; in particular, we focus on three of them, which have been implemented in `KPopTwistDB`:

#### Flat

This is *µ*(*I*_*i*_) = 1, i.e., no inertia-dependent scaling is performed.

#### Power

This is *µ*(*I*_*i*_) = (*I*_*i*_)^*q*^, *q* ≥ 0, i.e., the inertias are scaled according to some power *q*. This allows to implement schemes whereby the contribution to distance of each dimension is weighted by the amount of inertia associated with the dimension.

#### Thresholded power

The previous function can be generalised in order to include a threshold on the cumulative sum of rescaled inertias *C*_*i*_ across the first *i* dimensions, 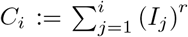, the rationale being that in certain situations we would only like to consider a limited number of dimensions, up for instance to some fraction of explained inertia, when we compute distances. Given an “internal” power *r* ≥ 0, an “external” power *s* ≥ 0, and a threshold 0 ≤ *t* ≤ 1, we would then generate the vector ***T*** defined by

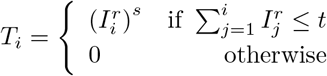

and return its normalised version as the result.

More details can be found on the `KPop` GitHub repository [35].

The *χ*^2^-distance traditionally used in correspondence analysis [69] can be recovered from the previous formulas by choosing Euclidean distances (*p* = 2) in Equation 3 and weighting dimensions by unscaled inertias, i.e. taking a Power metric with *q* = 1. These are the choices used throughout the paper. Note that the *χ*^2^distance is a Mahalanobis distance [69].

### KPop workflows

Based on the modular set of programs just introduced, one can implement arbitrarily complex workflows; some of them are illustrated in Figures 1, 2, 4, and 7. For each of these workflows, some steps can be omitted depending on the nature of the dataset being analysed; for instance, one would not typically perform the read selection of Figure 4 for environmental datasets, datasets obtained from bacterial isolates, or datasets obtained from targeted sequencing. A more detailed list of commands and options can be found online in the `KPop` GitHub repository [35].

### Comparison with MinHash-based methods

It should be noted that, strictly speaking, a direct comparison between `KPop` and MinHash-based methods such as `mash` [28] or `sourmash` [29] is not possible. That is because (see left-hand side of Figure 8) for each sequence or sample `KPop` produces an *embedding*, i.e., a point in a multi-dimensional space, which can be directly provided to classification methods as a numerical vector. On the other hand, for each sequence or sample MinHash-based methods produce a *sketch*, i.e., a representative set of hashes. The latter can be used to approximately compute distances between sequences or samples (right-hand side of Figure 8) but not be directly given as input to a classifier (red arrow on the right), as there is a very large number of possible MinHash *k*-mers at the values of *k* usually used by methods such as `mash` [28] or `sourmash` [29], and the dimension of the resulting vector would be given by the size of the intersection of all the hashes for all samples, which is usually very large.

This is why in this paper comparisons with MinHash-based methods were performed by using either built-in algorithms implemented by the method (as in the case of `sourmash tax`) or distance-based algorithms such as *K*-NN. The latter use the class of the *K* nearest neighbours in the training set to classify a test sequence (bottom right of Figure 8); as they only need the genomic distance to operate, an (indirect) comparison becomes possible. When using `sourmash tax`, we first built a database of “training” sketches with `sourmash index` and subsequently used it as input to `sourmash gather` to match the sketch derived for each test sequence with `sourmash sketch dna`; finally, `sourmash tax genome` was used to annotate the results of `sourmash gather` with information about the class or taxonomy. Distances were computed on all datasets using the command `mashtree --outmatrix` for `mash`, and with the command `sourmash sketch dna` followed by `sourmash compare *.sig --distance-matrix` for `sourmash`. All experiments running `sourmash` were performed with both *k* = 21 and *k* = 31 (the default).

## Author’s contributions

All authors contributed to this research and resulting manuscript. Their contributions are listed here in accordance with the Contributor Roles Taxonomy (CRediT). Xavier Didelot: Methodology, Investigation, Writing - original draft, Writing - review & editing. Paolo Ribeca: Conceptualization, Methodology, Software, Investigation, Writing - original draft, Writing - review & editing.

## Acknowledgements

This study was partly funded by the National Institute for Health Research (NIHR) Health Protection Research Unit in Genomics and Enabling Data.

The authors acknowledge the Research/Scientific Computing teams at The James Hutton Institute and NIAB for providing computational resources and technical support for the UK’s Crop Diversity Bioinformatics HPC (BBSRC grant BB/S019669/1), use of which has contributed to the results reported within this paper. The invaluable assistance constantly provided by Dr Iain Milne is especially noted and appreciated.

Finally, the authors are indebted to Dr Ryan Morrison for his help with improving online documentation and setting up the KPop conda channel.

## Availability of data and materials

All code is open source and available under the terms of the GNU General Public License v3.0 on the GitHub repository of KPop at https://github.com/PaoloRibeca/KPop.

The tuberculosis and SARS-CoV-2 simulated datasets can be reproduced using code provided in the test directory. The accession numbers of the *Mycobacterium* genomes used for the classifier application are also given in the test directory.

The SARS-CoV-2 Pangolin lineage dataset is available from https://raw.githubusercontent.com/cov-lineages/pango-designation/master/lineages.csv and the corresponding genome sequencing are available from GISAID at https://www.gisaid.org/.

## Supplementary Figures

**Supplementary Figure 1.**
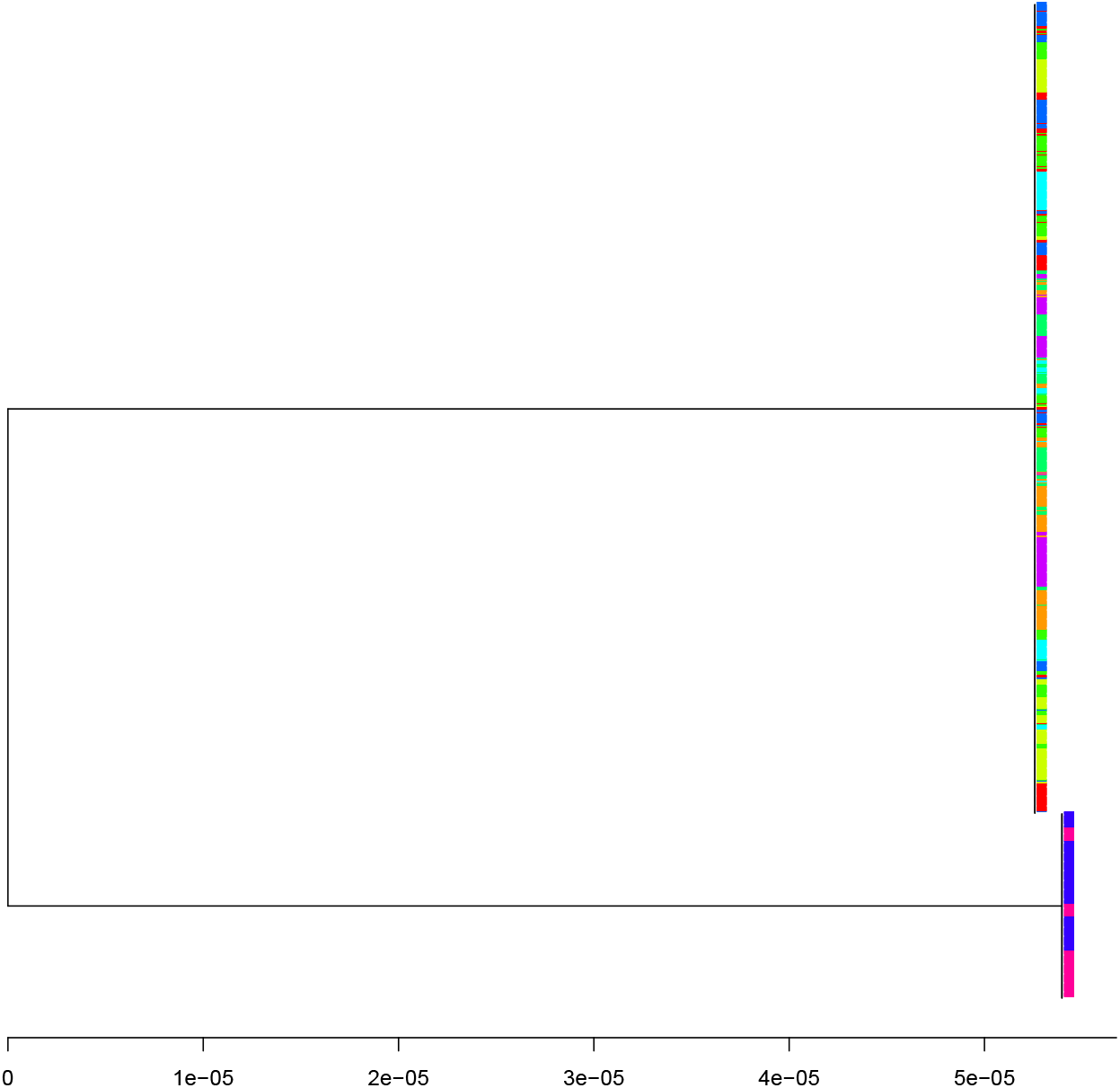
Tree produced by MashTree on the simulated tuberculosis dataset. The leaves of the tree are shown using unique colours for each of the ten lineages as in Figure 3.

**Supplementary Figure 2.**
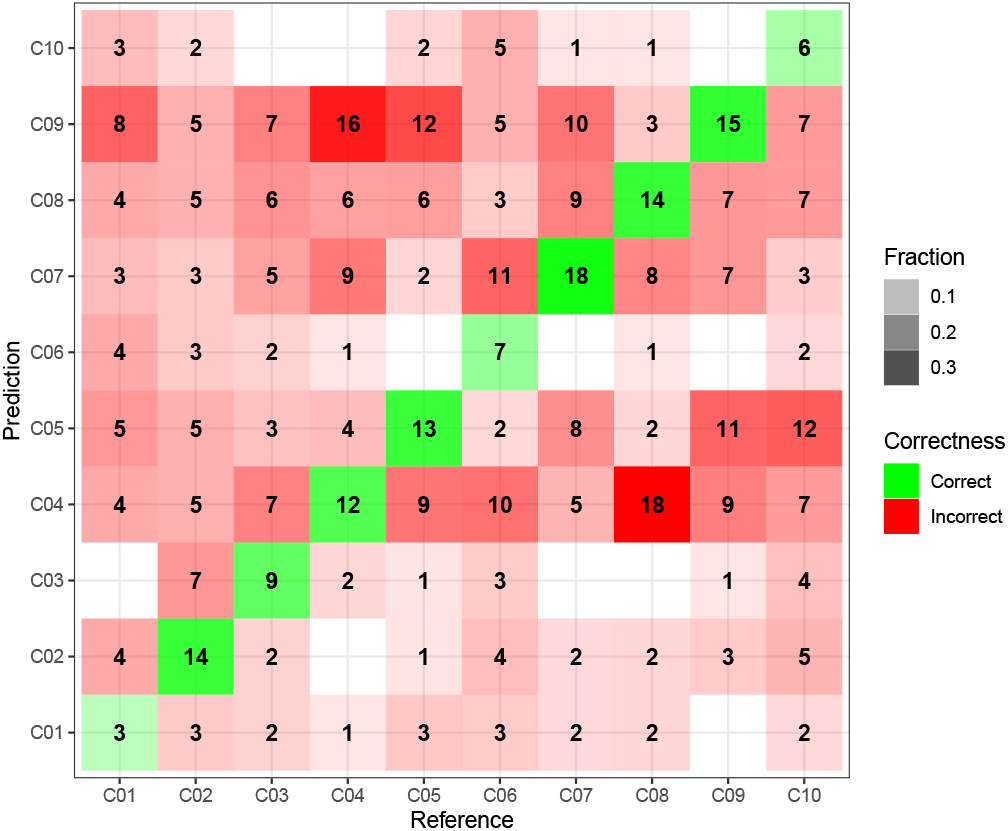
Confusion matrix for the results of sourmash tax on the simulated tuberculosis dataset at *k* = 31. The ten classes in the figure (named C1 to C10) correspond to the ten lineages in Figure 3.

**Supplementary Figure 3.**
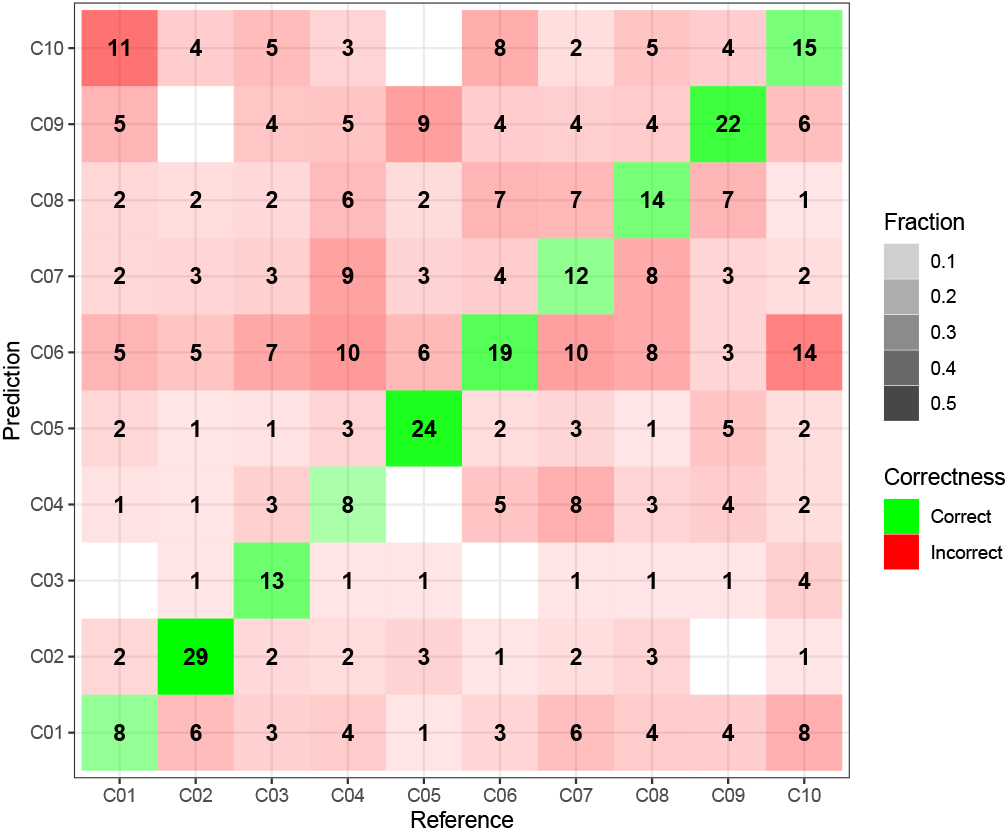
Confusion matrix for the results of sourmash tax on the simulated tuberculosis dataset at *k* = 21. The ten classes in the figure (named C1 to C10) correspond to the ten lineages in Figure 3.

**Supplementary Figure 4.**
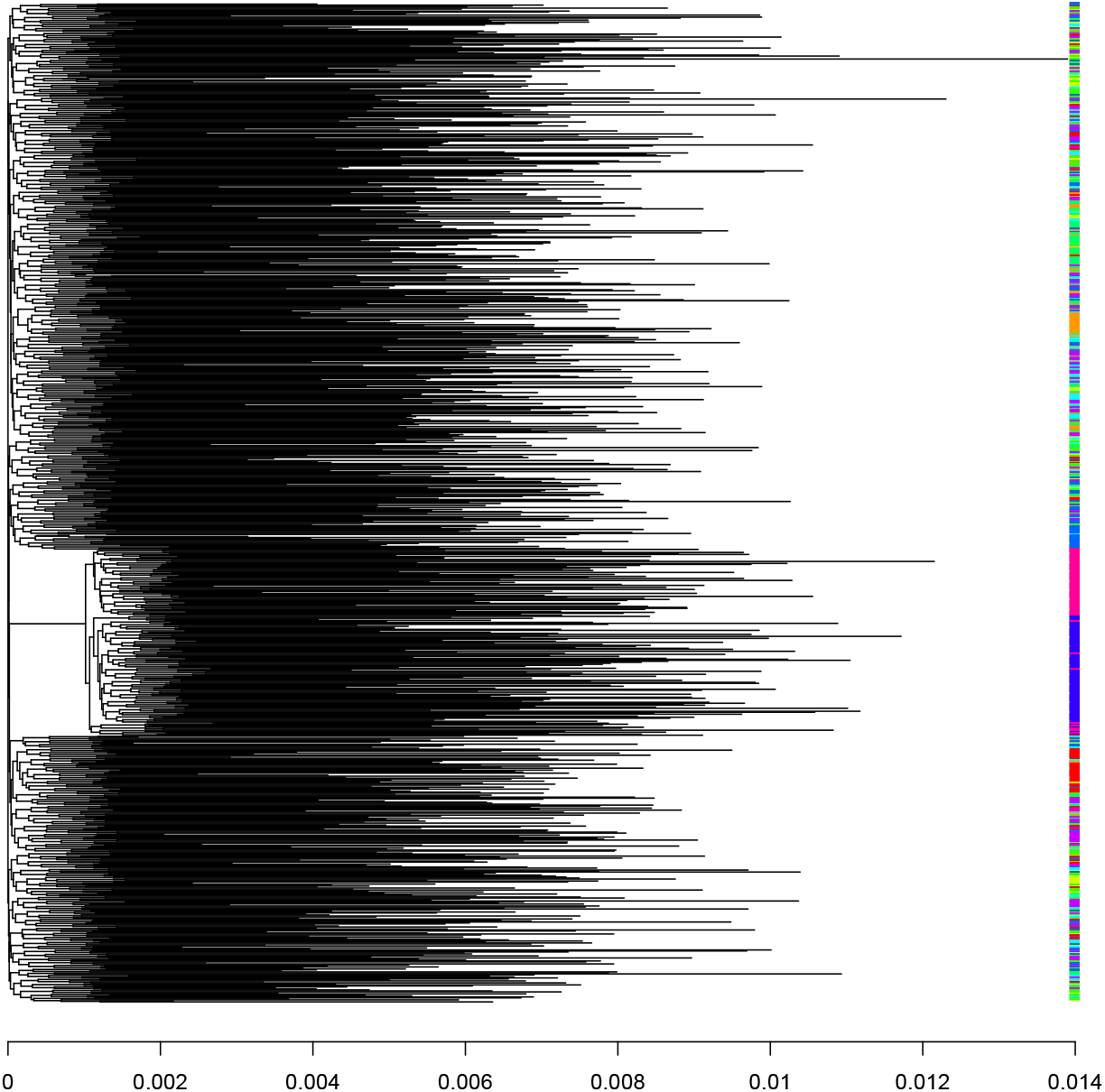
Neighbour-joining tree based on the distances estimated by sourmash on the simulated tuberculosis dataset. The leaves of the tree are shown using unique colours for each of the ten lineages as in Figure 3.

**Supplementary Figure 5.**
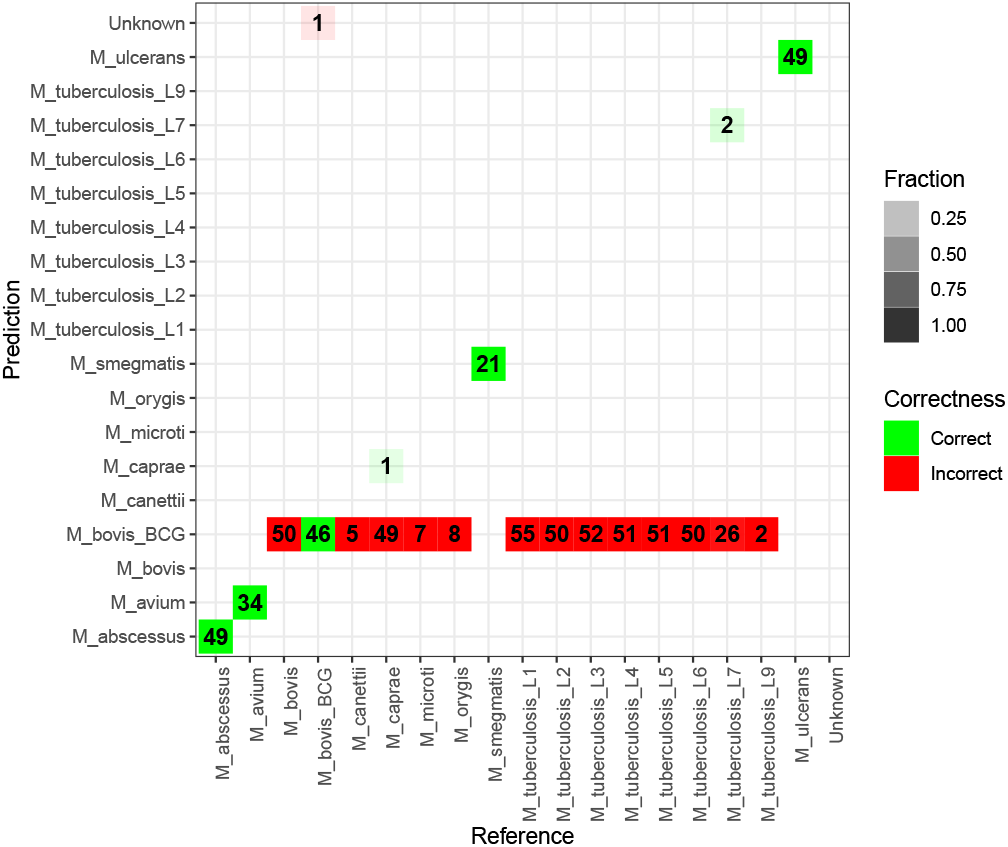
Confusion matrix for the results of sourmash tax on the Mycobacterium sequencing dataset at *k* = 31.

**Supplementary Figure 6.**
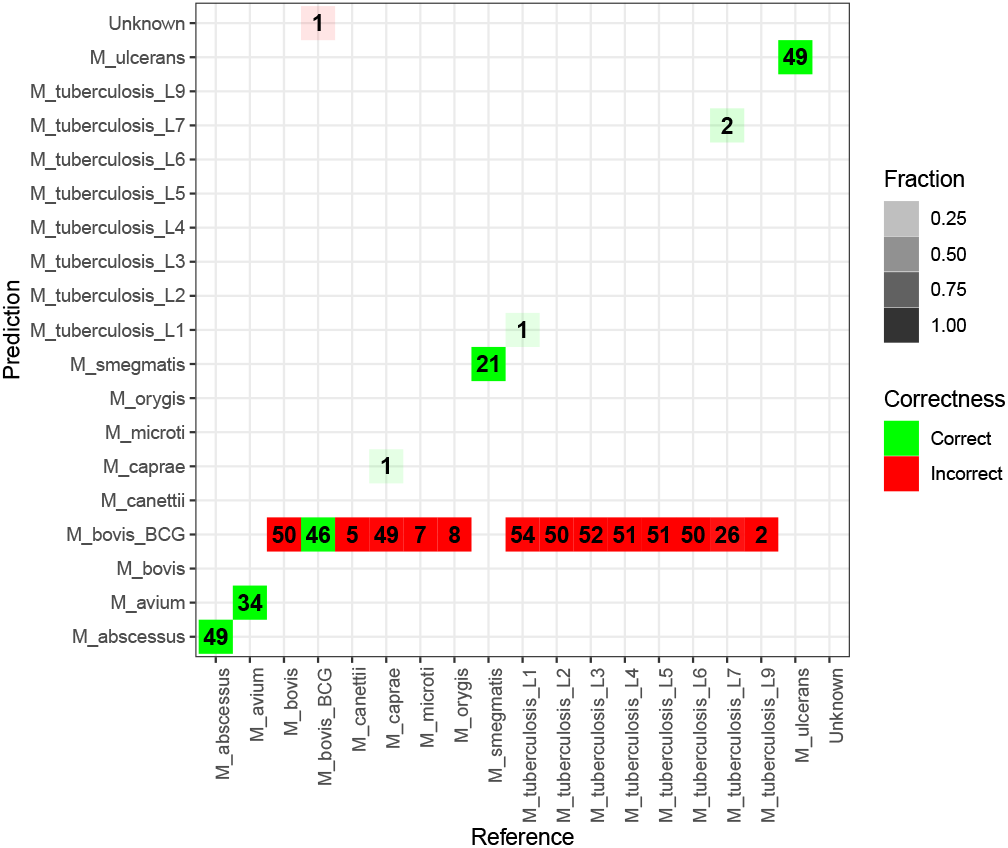
Confusion matrix for the results of sourmash tax on the Mycobacterium sequencing dataset at *k* = 21.

**Supplementary Figure 7.**
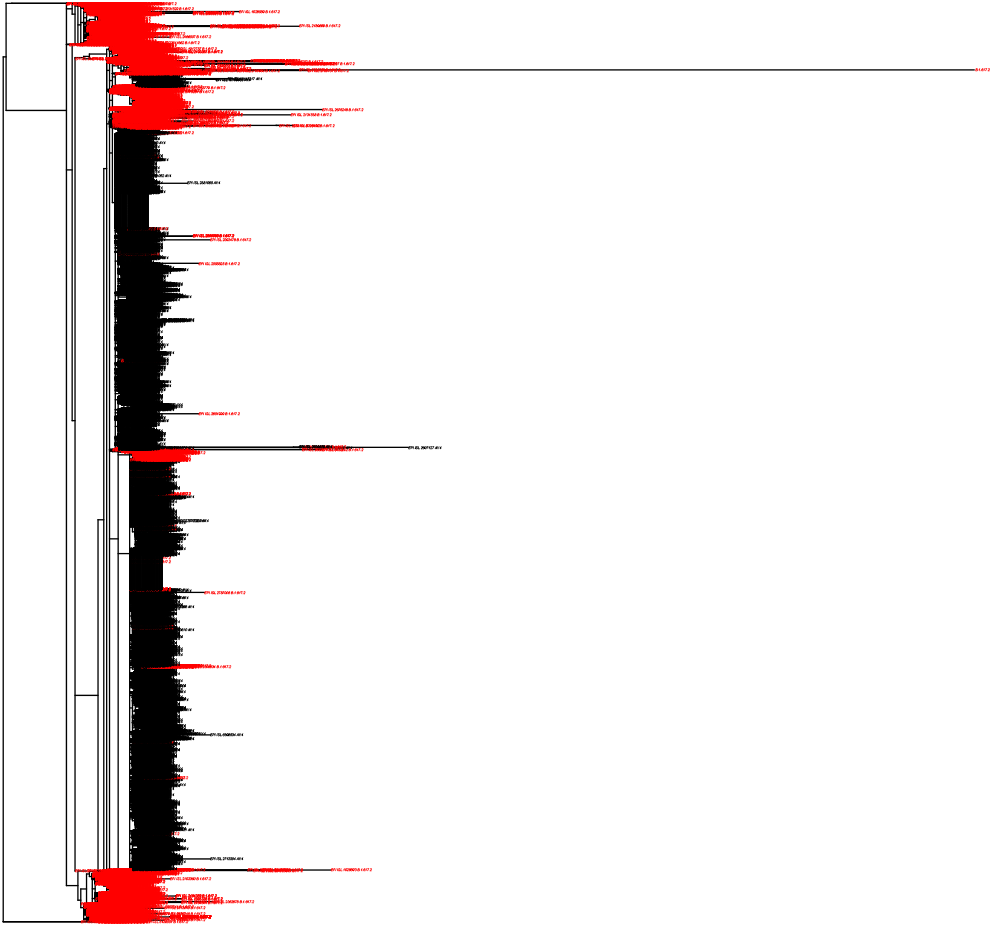
Clonal genealogy used by SimBac when simulating the dataset of 100 genomes from a recombinant population.

**Supplementary Figure 8.**
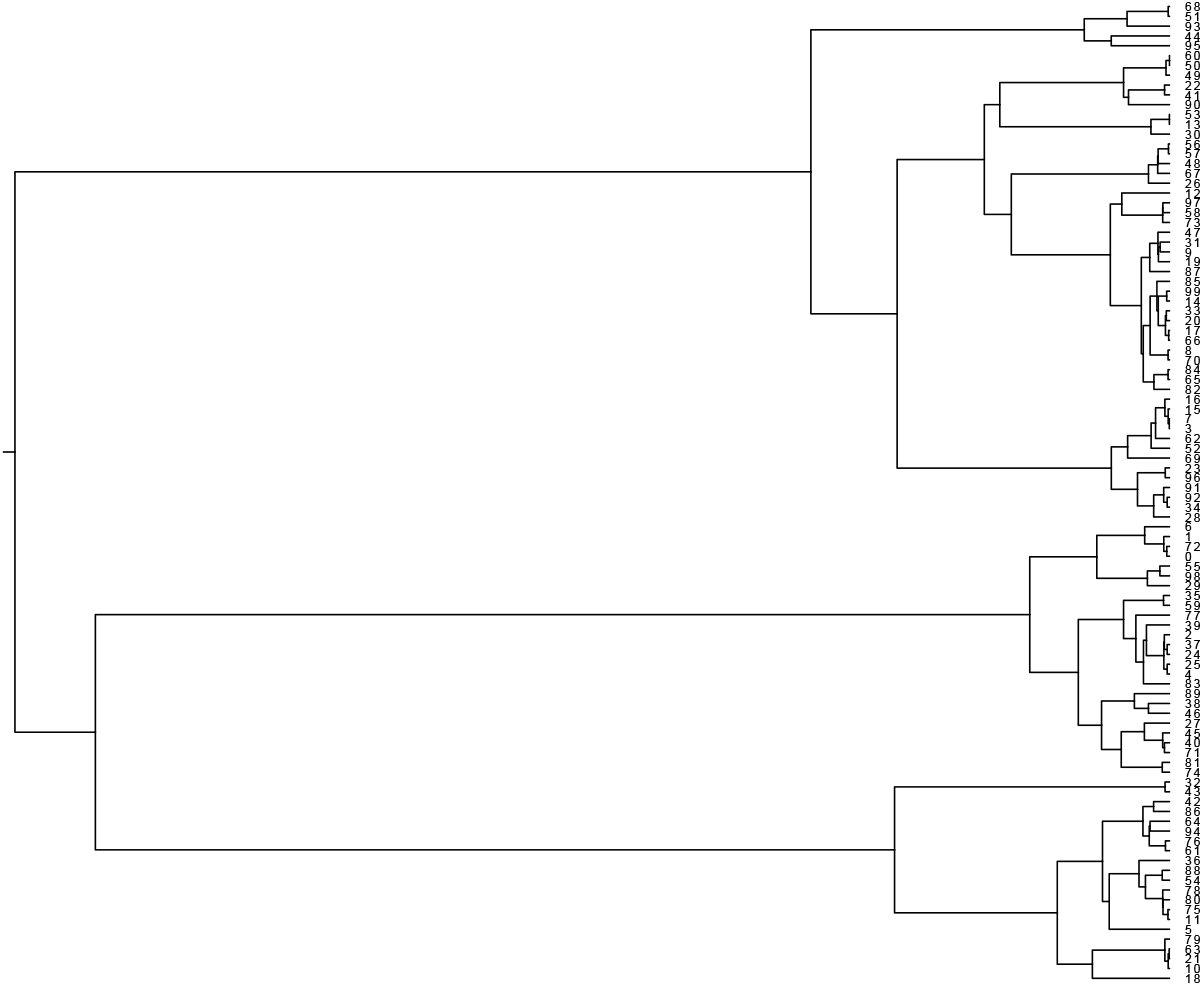
Tree produced by Usher on the SARS-CoV-2 dataset, showing only in red the genomes from lineage B.1.617.2 and in black the genomes from lineage AY.4 (alias of B.1.617.2.4).

## References

1. Didelot, X., Bowden, R., Wilson, D.J., Peto, T.E.A., Crook, D.W.: Transforming clinical microbiology with bacterial genome sequencing. Nat. Rev. Genet. 13, 601–612 (2012). doi:10.1038/nrg3226

2. Loman, N.J., Pallen, M.J.: Twenty years of bacterial genome sequencing. Nat. Rev. Microbiol. 13, 787–94 (2015). doi:10.1038/nrmicro3565

3. Goodwin, S., McPherson, J.D., McCombie, W.R.: Coming of age: ten years of next-generation sequencing technologies. Nat. Rev. Genet. 17(6), 333–351 (2016). doi:10.1038/nrg.2016.49

4. Gardy, J.L., Loman, N.J.: Towards a genomics-informed, real-time, global pathogen surveillance system. Nat. Rev. Genet. 19, 9–20 (2018). doi:10.1038/nrg.2017.88

5. Benson, D.A., Cavanaugh, M., Clark, K., Karsch-Mizrachi, I., Lipman, D.J., Ostell, J., Sayers, E.W.: GenBank. Nucleic Acids Res. 45(D1), 37–42 (2017). doi:10.1093/nar/gkw1070

6. Jolley, K.A., Bray, J.E., Maiden, M.C.J.: Open-access bacterial population genomics: BIGSdb software, the PubMLST.org website and their applications [version 1; referees: 2 approved]. Wellcome Open Res. 3(0), 1–20 (2018). doi:10.12688/wellcomeopenres.14826.1

7. Zhou, Z., Alikhan, N.F., Mohamed, K., Fan, Y., Achtman, M.: The EnteroBase user’s guide, with case studies on Salmonella transmissions, Yersinia pestis phylogeny, and Escherichia core genomic diversity. Genome Res. 30(1), 138–152 (2020). doi:10.1101/gr.251678.119

8. Li, H., Durbin, R.: Fast and accurate short read alignment with Burrows?Wheeler transform. Bioinformatics 25(14), 1754–1760 (2009)

9. Langmead, B., Salzberg, S.L.: Fast gapped-read alignment with Bowtie 2. Nat. Methods 9(4), 357–359 (2012). doi:10.1038/nmeth.1923

10. Marco-Sola, S., Sammeth, M., Guigó, R., Ribeca, P.: The GEM mapper: fast, accurate and versatile alignment by filtration. Nature methods 9(12), 1185–1188 (2012)

11. Garrison, E., Marth, G.: Haplotype-based variant detection from short-read sequencing. arXiv preprint arXiv:1207.3907 (2012)

12. Zerbino, D.R., Birney, E.: Velvet: Algorithms for de novo short read assembly using de Bruijn graphs. Genome Res. 18, 821–829 (2008). doi:10.1101/gr.074492.107

13. Bankevich, A., Nurk, S., Antipov, D., Gurevich, A.a., Dvorkin, M., Kulikov, A.S., Lesin, V.M., Nikolenko, S.I., Pham, S., Prjibelski, A.D., Pyshkin, A.V., Sirotkin, A.V., Vyahhi, N., Tesler, G., Alekseyev, M.a., Pevzner, P.a.: SPAdes: A New Genome Assembly Algorithm and Its Applications to Single-Cell Sequencing. J. Comput. Biol. 19(5), 455–477 (2012). doi:10.1089/cmb.2012.0021

14. Nagarajan, N., Pop, M.: Sequence assembly demystified. Nat. Rev. Genet. 14(3), 157–67 (2013). doi:10.1038/nrg3367

15. Vernikos, G., Medini, D., Riley, D.R., Tettelin, H.: Ten years of pan-genome analyses. Curr. Opin. Microbiol. 23, 148–154 (2015). doi:10.1016/j.mib.2014.11.016

16. Darling, A.E., Mau, B., Perna, N.T.: progressiveMauve: Multiple Genome Alignment with Gene Gain, Loss and Rearrangement. PLoS One 5(6), 11147 (2010). doi:10.1371/journal.pone.0011147

17. Angiuoli, S.V., Salzberg, S.L.: Mugsy: fast multiple alignment of closely related whole genomes. Bioinformatics 27(3), 334–42 (2011)

18. Maiden, M.C.J., Jansen van Rensburg, M.J., Bray, J.E., Earle, S.G., Ford, S.a., Jolley, K.a., McCarthy, N.D.: MLST revisited: the gene-by-gene approach to bacterial genomics. Nat. Rev. Microbiol. 11(10), 728–36 (2013). doi:10.1038/nrmicro3093

19. Kurtz, S., Phillippy, A., Delcher, A.L., Smoot, M., Shumway, M., Antonescu, C., Salzberg, S.L.: Versatile and open software for comparing large genomes. Genome Biol. 5(2), 12 (2004). doi:10.1186/gb-2004-5-2-r12

20. Didelot, X., Darling, A., Falush, D.: Inferring genomic flux in bacteria. Genome Res. 19(2), 306–317 (2008). doi:10.1101/gr.082263.108

21. Sims, G.E., Kim, S.-H.: Whole-genome phylogeny of Escherichia coli/Shigella group by feature frequency profiles (FFPs). Proc Natl Acad Sci USA 108(20), 8329–34 (2011). doi:10.1073/pnas.1105168108

22. Sheppard, S.K., Didelot, X., Meric, G., Torralbo, A., Jolley, K.A., Kelly, D.J., Bentley, S.D., Maiden, M.C.J., Parkhill, J., Falush, D.: Genome-wide association study identifies vitamin B5 biosynthesis as a host specificity factor in Campylobacter. Proc Natl Acad Sci USA 110(29), 11923–7 (2013). doi:10.5061/dryad.28n35.

23. Méric, G., Mageiros, L., Pensar, J., Laabei, M., Yahara, K., Pascoe, B., Kittiwan, N., Tadee, P., Post, V., Lamble, S., Bowden, R., Bray, J.E., Morgenstern, M., Jolley, K.A., Maiden, M.C.J., Feil, E.J., Didelot, X., Miragaia, M., de Lencastre, H., Moriarty, T.F., Rohde, H., Massey, R., Mack, D., Corander, J., Sheppard, S.K.: Disease-associated genotypes of the commensal skin bacterium Staphylococcus epidermidis. Nat. Commun. 9(1), 5034 (2018). doi:10.1038/s41467-018-07368-7

24. Bradley, P., Gordon, N.C., Walker, T.M., Dunn, L., Heys, S., Huang, B., Earle, S., Pankhurst, L.J., Anson, L., de Cesare, M., Piazza, P., Votintseva, A.A., Golubchik, T., Wilson, D.J., Wyllie, D.H., Diel, R., Niemann, S., Feuerriegel, S., Kohl, T.A., Ismail, N., Omar, S.V., Smith, E.G., Buck, D., McVean, G., Walker, A.S., Peto, T., Crook, D.W., Iqbal, Z.: Rapid antibiotic resistance predictions from genome sequence data for S. aureus and M. tuberculosis. Nat Comm 6, 018564 (2015). doi:10.1101/018564. arXiv:1011.1669v3

25. Alneberg, J., Bjarnason, B.S., De Bruijn, I., Schirmer, M., Quick, J., Ijaz, U.Z., Lahti, L., Loman, N.J., Andersson, A.F., Quince, C.: Binning metagenomic contigs by coverage and composition. Nature methods 11(11), 1144–1146 (2014)

26. Laczny, C.C., Pinel, N., Vlassis, N., Wilmes, P.: Alignment-free visualization of metagenomic data by nonlinear dimension reduction. Scientific reports 4(1), 1–12 (2014)

27. Pan, S., Zhu, C., Zhao, X.-M., Coelho, L.P.: A deep siamese neural network improves metagenome-assembled genomes in microbiome datasets across different environments. Nature Communications 13(1), 1–12 (2022)

28. Ondov, B.D., Treangen, T.J., Melsted, P., Mallonee, A.B., Bergman, N.H., Koren, S., Phillippy, A.M.: Mash: Fast genome and metagenome distance estimation using MinHash. Genome Biol. 17(1), 1–14 (2016). doi:10.1186/s13059-016-0997-x

29. Brown, C.T., Irber, L.: sourmash: a library for minhash sketching of dna. Journal of open source software 1(5), 27 (2016)

30. Lees, J.A., Harris, S.R., Tonkin-Hill, G., Gladstone, R.A., Lo, S.W., Weiser, J.N., Corander, J., Bentley, S.D., Croucher, N.J.: Fast and flexible bacterial genomic epidemiology with PopPUNK. Genome Res. 29(2), 304–316 (2019). doi:10.1101/gr.241455.118

31. Belbasi, M., Blanca, A., Harris, R.S., Koslicki, D., Medvedev, P.: The minimizer Jaccard estimator is biased and inconsistent. Bioinformatics 38(Supplement 1), 169–176 (2022). doi:10.1093/bioinformatics/btac244. Accessed 2022-07-01

32. Ripley, B.: Classification and regression trees [R package tree version 1. 0-42]. Comprehensive R Archive Network (CRAN) (2022). https://cran.r-project.org/web/packages/tree/index.html

33. Liaw, A., Wiener, M.: Classification and Regression by randomForest. R News 2(3), 18–22 (2002)

34. Didelot, X., Wilson, D.J.: ClonalFrameML: efficient inference of recombination in whole bacterial genomes. PLoS computational biology 11(2), 1004041 (2015)

35. KPop source code and distributions. https://github.com/PaoloRibeca/KPop. Accessed: 2022-05-30

36. Blackwell, G.A., Hunt, M., Malone, K.M., Lima, L., Horesh, G., Alako, B.T., Thomson, N.R., Iqbal, Z.: Exploring bacterial diversity via a curated and searchable snapshot of archived dna sequences. PLoS biology 19(11), 3001421 (2021)

37. Sheppard, S.K., Didelot, X., Jolley, K.A., Darling, A.E., Pascoe, B., Meric, G., Kelly, D.J., Cody, A., Colles, F.M., Strachan, N.J.C., Ogden, I.D., Forbes, K., French, N.P., Carter, P., Miller, W.G., McCarthy, N.D., Owen, R., Litrup, E., Egholm, M., Affourtit, J.P., Bentley, S.D., Parkhill, J., Maiden, M.C.J., Falush, D.: Progressive genome-wide introgression in agricultural Campylobacter coli. Molecular Ecology 22, 1051–1064 (2013). doi:10.1111/mec.12162

38. Menardo, F., Duchêne, S., Brites, D., Gagneux, S.: The molecular clock of Mycobacterium tuberculosis. PLoS Pathog. 15(9), 1–24 (2019). doi:10.1371/journal.ppat.1008067

39. Cole, S., Brosch, R., Parkhill, J., Garnier, T., Churcher, C., Harris, D., Gordon, S.V., Eiglmeier, K., Gas, S., Barry, C.E.r., Others: Deciphering the biology of Mycobacterium tuberculosis from the complete genome sequence. Nature 396, 537–544 (1998)

40. Huang, W., Li, L., Myers, J.R., Marth, G.T.: ART: A next-generation sequencing read simulator. Bioinformatics 28(4), 593–594 (2012). doi:10.1093/bioinformatics/btr708

41. Cover, T., Hart, P.: Nearest neighbor pattern classification. IEEE transactions on information theory 13(1), 21–27 (1967)

42. Katz, L.S., Griswold, T., Morrison, S.S., Caravas, J.A., Zhang, S., Bakker, H.C.d., Deng, X., Carleton, H.A.: Mashtree: a rapid comparison of whole genome sequence files. Journal of Open Source Software 4(44), 1762 (2019). doi:10.21105/joss.01762. Accessed 2022-06-07

43. The NCBI Short Read Archive. https://www.ncbi.nlm.nih.gov/sra. Accessed: 2022-05-30

44. Phelan, J.E., O’Sullivan, D.M., Machado, D., Ramos, J., Oppong, Y.E.A., Campino, S., O’Grady, J., McNerney, R., Hibberd, M.L., Viveiros, M., Huggett, J.F., Clark, T.G.: Integrating informatics tools and portable sequencing technology for rapid detection of resistance to anti-tuberculous drugs. Genome Med. 11(1), 1–7 (2019). doi:10.1186/s13073-019-0650-x

45. Brites, D., Loiseau, C., Menardo, F., Borrell, S., Boniotti, M.B., Warren, R., Dippenaar, A., Parsons, S.D.C., Beisel, C., Behr, M.A., Fyfe, J.A., Coscolla, M., Gagneux, S.: A new phylogenetic framework for the animal-adapted mycobacterium tuberculosis complex. Front. Microbiol. 9(NOV), 1–14 (2018). doi:10.3389/fmicb.2018.02820

46. Trim Galore. https://www.bioinformatics.babraham.ac.uk/projects/trim_galore. Accessed: 2022-05-30

47. Mago?c, T., Salzberg, S.L.: FLASH: fast length adjustment of short reads to improve genome assemblies. Bioinformatics 27(21), 2957–2963 (2011)

48. Eldholm, V., Balloux, F.: Antimicrobial resistance in mycobacterium tuberculosis: the odd one out. Trends in microbiology 24(8), 637–648 (2016)

49. Brown, T., Didelot, X., Wilson, D.J., De Maio, N.: SimBac: simulation of whole bacterial genomes with homologous recombination. Microbial genomics 2(1) (2016)

50. Zhou, Z., Charlesworth, J., Achtman, M.: Accurate reconstruction of bacterial pan-and core genomes with PEPPAN. Genome research 30(11), 1667–1679 (2020)

51. Wu, F., Zhao, S., Yu, B., Chen, Y.-M., Wang, W., Song, Z.-G., Hu, Y., Tao, Z.-W., Tian, J.-H., Pei, Y.-Y., Yuan, M.-L., Zhang, Y.-L., Dai, F.-H., Liu, Y., Wang, Q.-M., Zheng, J.-J., Xu, L., Holmes, E.C., Zhang, Y.-Z.: A new coronavirus associated with human respiratory disease in China. Nature 579(7798), 265–269 (2020). doi:10.1038/s41586-020-2008-3

52. van Dorp, L., Acman, M., Richard, D., Shaw, L.P., Ford, C.E., Ormond, L., Owen, C.J., Pang, J., Tan, C.C.S., Boshier, F.A.T., Ortiz, A.T., Balloux, F.: Emergence of genomic diversity and recurrent mutations in SARS-CoV-2. Infect. Genet. Evol. 83(April), 104351 (2020). doi:10.1016/j.meegid.2020.104351

53. O’Toole, Á., Scher, E., Underwood, A., Jackson, B., Hill, V., McCrone, J.T., Colquhoun, R., Ruis, C., Abu-Dahab, K., Taylor, B., Yeats, C., du Plessis, L., Maloney, D., Medd, N., Attwood, S.W., Aanensen, D.M., Holmes, E.C., Pybus, O.G., Rambaut, A.: Assignment of epidemiological lineages in an emerging pandemic using the pangolin tool. Virus Evol. 7(2), 1–9 (2021). doi:10.1093/ve/veab064

54. Pangolin Lineage Dataset. https://raw.githubusercontent.com/cov-lineages/pango-designation/master/lineages.csv. Accessed: 2022-05-30

55. Turakhia, Y., Thornlow, B., Hinrichs, A.S., De Maio, N., Gozashti, L., Lanfear, R., Haussler, D., Corbett-Detig, R.: Ultrafast Sample placement on Existing tRees (UShER) enables real-time phylogenetics for the SARS-CoV-2 pandemic. Nature Genetics 53(6), 809–816 (2021)

56. Sunagawa, S., Coelho, L.P., Chaffron, S., Kultima, J.R., Labadie, K., Salazar, G., Djahanschiri, B., Zeller, G., Mende, D.R., Alberti, A., et al.: Structure and function of the global ocean microbiome. Science 348(6237), 1261359 (2015)

57. Delmont, T.O., Quince, C., Shaiber, A., Esen, Ö.C., Lee, S.T., Rappé, M.S., MacLellan, S.L., Lücker, S., Eren, A.M.: Nitrogen-fixing populations of Planctomycetes and Proteobacteria are abundant in surface ocean metagenomes. Nature Microbiology (2018). doi:10.1038/s41564-018-0176-9

58. Dutta, A., Pellow, D., Shamir, R.: Parameterized syncmer schemes improve long-read mapping. PLOS Computational Biology 18(10), 1–19 (2022). doi:10.1371/journal.pcbi.1010638

59. Kille, B., Garrison, E., Treangen, T.J., Phillippy, A.M.: Minmers are a generalization of minimizers that enable unbiased local Jaccard estimation. Bioinformatics 39(9), 512 (2023). doi:10.1093/bioinformatics/btad512. https://academic.oup.com/bioinformatics/article-pdf/39/9/btad512/51648656/btad512.pdf

60. Shaw, J., Yu, Y.W.: Metagenome profiling and containment estimation through abundance-corrected k-mer sketching with sylph. bioRxiv (2024). doi:10.1101/2023.11.20.567879. https://www.biorxiv.org/content/early/2024/01/22/2023.11.20.567879.full.pdf

61. Wang, J., Yi, X., Guo, R., Jin, H., Xu, P., Li, S., Wang, X., Guo, X., Li, C., Xu, X., et al.: Milvus: A purpose-built vector data management system. In: Proceedings of the 2021 International Conference on Management of Data, pp. 2614–2627 (2021)

62. Johnson, J., Douze, M., Jegou, H.: Billion-scale similarity search with gpus. IEEE Transactions on Big Data 7(03), 535–547 (2021). doi:10.1109/TBDATA.2019.2921572

63. Kapli, P., Yang, Z., Telford, M.J.: Phylogenetic tree building in the genomic age. Nat. Rev. Genet. 21(July) (2020). doi:10.1038/s41576-020-0233-0

64. The OCaml programming language. https://ocaml.org. Accessed: 2022-05-30

65. R Core Team: R: A Language and Environment for Statistical Computing. R Foundation for Statistical Computing, Vienna, Austria (2021). R Foundation for Statistical Computing. https://www.R-project.org/

66. Benzécri, J.-P.: Correspondence Analysis Handbook. CRC Press LLC, ??? (1992)

67. Nenadic, O., Greenacre, M.: Correspondence analysis in r, with two- and three-dimensional graphics: The ca package. Journal of Statistical Software 20(3), 1–13 (2007)

68. Ghandi, M., Lee, D., Mohammad-Noori, M., Beer, M.A.: Enhanced regulatory sequence prediction using gapped k-mer features. PLoS computational biology 10(7), 1003711 (2014)

69. Greenacre, M.: Correspondence Analysis in Practice. Chapman and Hall/CRC, ??? (2007)

70. van Dam, A., Dekker, M., Morales-Castilla, I., Rodríguez, M., Wichmann, D., Baudena, M.: Correspondence analysis, spectral clustering and graph embedding: applications to ecology and economic complexity. Sci. Rep. 11(1), 8926 (2021). doi:10.1038/s41598-021-87971-9

